# Decay chemicals activate a pH-sensing pathway to initiate a persistent state

**DOI:** 10.64898/2026.09.16.751831

**Authors:** Grace M. Padilla, Matthew L. White, Ming Tak Ngan, John Jefferson, Jennifer Swindlehurst Chan, Kathleen A. Martin, Matthew Lovett-Barron

## Abstract

Animals use multiple sensory systems to identify signs of danger, including dead conspecifics. Fish can identify decaying conspecifics from chemosensory cues, which prompt a long-lasting avoidance; the neural basis of this persistent chemosensory-driven state is not understood. Here, we describe a persistent alarm state in larval zebrafish, where brief exposure to the polyamine decay product, cadaverine, initiates a minutes-long increase in heart rate and suppression of visually-evoked swimming. Cadaverine activates both olfactory and trigeminal neurons, with persistently-active cells in the forebrain and dorsolateral hindbrain. However, we find that cadaverine’s persistent effects on hindbrain activity, heart rate, and locomotor suppression are not mediated by olfaction. Instead, they are mediated through trigeminal chemosensation of the basic pH of concentrated cadaverine in solution. Knockout of TrpA1 channels, which are expressed in trigeminal neurons and associated with chemical nociception across species, prevents the persistent suppression of movement from brief exposure to cadaverine. pH-responsive trigeminal neurons project into the dorsolateral hindbrain to mediate persistent activity and long-lasting locomotor suppression through local inhibition. These results demonstrate that non-olfactory chemosensation mediates persistent behavioral responses to death and decay in zebrafish, illustrating how diverse chemosensory systems detect ethologically-relevant cues to drive adaptive behavior.

## Introduction

Animals have evolved a wide variety of sensory systems, allowing them to detect and respond to features of their environment that are relevant to their survival^1,2^. For instance, it is critically important for animals to detect stimuli that threaten their life, to engage in behaviors that maximize their chances of outlasting such threats and avoiding harm. Detecting threats can lead to a rapid avoidance response, but can also elicit long-timescale changes in behavior^3,4^ that outlast the sensory event itself. Such persistent states are manifested in altered neural activity and behavioral expression^4–13^, and allow animals to respond to threat-related sensory cues by adjusting their behaviors on longer timescales.

Among the many sensory cues that can indicate a threat, exposure to dead or decaying conspecifics can be a potent sign of danger^3,14–16^, not only from the potential predators that may have killed the conspecific, but also because decaying organic matter can spread disease and pathogens^17^. Many species change their behavior upon exposure to dead conspecifics, in response to a range of sensory cues^18–22^. Chemical indicators of decay, such as the bacterial products cadaverine and putrescine^23–25^, are sufficient to elicit strong aversion through activation of olfactory trace amine associated receptors (TAARs)^23,26–28^. TAARs are highly conserved across vertebrates, suggesting common mechanisms for sensation of tissue decay.

Alarm and avoidance behaviors in response to injury, death, or decay-related chemicals have been studied in zebrafish, where olfactory-mediated avoidance to the polyamine cadaverine is detected by a singular receptor in the zebrafish olfactory epithelium, TAAR13c (ref. ^23^). Despite this well-established olfactory mechanism for detecting cadaverine, there is growing evidence that polyamines can be detected through other receptors besides TAARs^26,28,29^. Zebrafish and other vertebrates possess a range of mechanisms for chemosensation^30–32^, suggesting other chemosensory modalities may be involved in the behavioral responses to decay-related chemicals.

Here we describe how brief exposure to high concentrations of the decay-related chemical cadaverine evokes a long-lasting state characterized by an elevated heart rate and suppressed locomotion in larval zebrafish. These behavioral and physiological measures are correlated with the activity of neurons in olfactory-related regions of the forebrain, as well as a population of neurons in the dorsolateral hindbrain. Surprisingly, we found that ablation of olfactory sensory neurons did not impact cadaverine’s persistent effects on heart rate, locomotor suppression, or hindbrain activity. Instead, we find that the persistent cadaverine-evoked state is mediated by alternative chemosensory pathways – including pH-sensitive neurons in the trigeminal ganglia – which receive sensory input from the face and whose axons project into the hindbrain regions where we observed persistent activity. Cadaverine, and naturally-decaying fish, have a basic pH and activate trigeminal neurons via the TrpA1b channel. This pH-sensitive hindbrain circuit drives local GABAergic inhibitory neurons, whose activation suppresses movement independent of heart rate. Our results demonstrate that, in addition to classical olfactory-mediated mechanisms of cadaverine avoidance, high-concentrations of cadaverine and other products of decay can recruit the trigeminal chemical nociception system through changes in pH, to produce a persistent internal state underlying adaptive behavior.

## Results

### Brief exposure to cadaverine induces persistent changes in physiology and behavior

We first exposed zebrafish larvae to cadaverine, a well-characterized decay-related odor known to cause avoidance behaviors in adults and larvae^23,33^. We measured the persistent behavioral and physiological effects of brief cadaverine exposure, by recording from agarose-tethered behaving zebrafish larvae (7-9 days post-fertilization) during and after one minute of cadaverine exposure (see **Methods**). We used this head-tethered preparation to enable precise stimulus control and high-resolution behavioral quantification in a configuration conducive to neuronal imaging (**Figure 1A**). We removed agarose around the front of their face and the tail, to allow odor detection and tail movements (**Figures 1B, S1A**). We tracked tail movements over a 25-minute video recording and analyzed eye saccades and heart rate from recorded videos (see **Methods, Figure 1B, S1A-B**), where we measured the baseline heart rate at 3.29±0.07 Hz (mean±sem, N=57; **Figure S1B**). We exposed tethered fish to a constant flow of standard embryo/larvae medium (E3; see **Methods**) for 5 minutes, then switched to a cadaverine solution for one minute before switching back to E3 for the remainder of the recording. A vacuum pipette provided constant suction to allow for rapid water exchange in the behavioral chamber, and to ensure that chemical delivery is not associated with abrupt changes in flow rate or water level.

**Figure 1.**
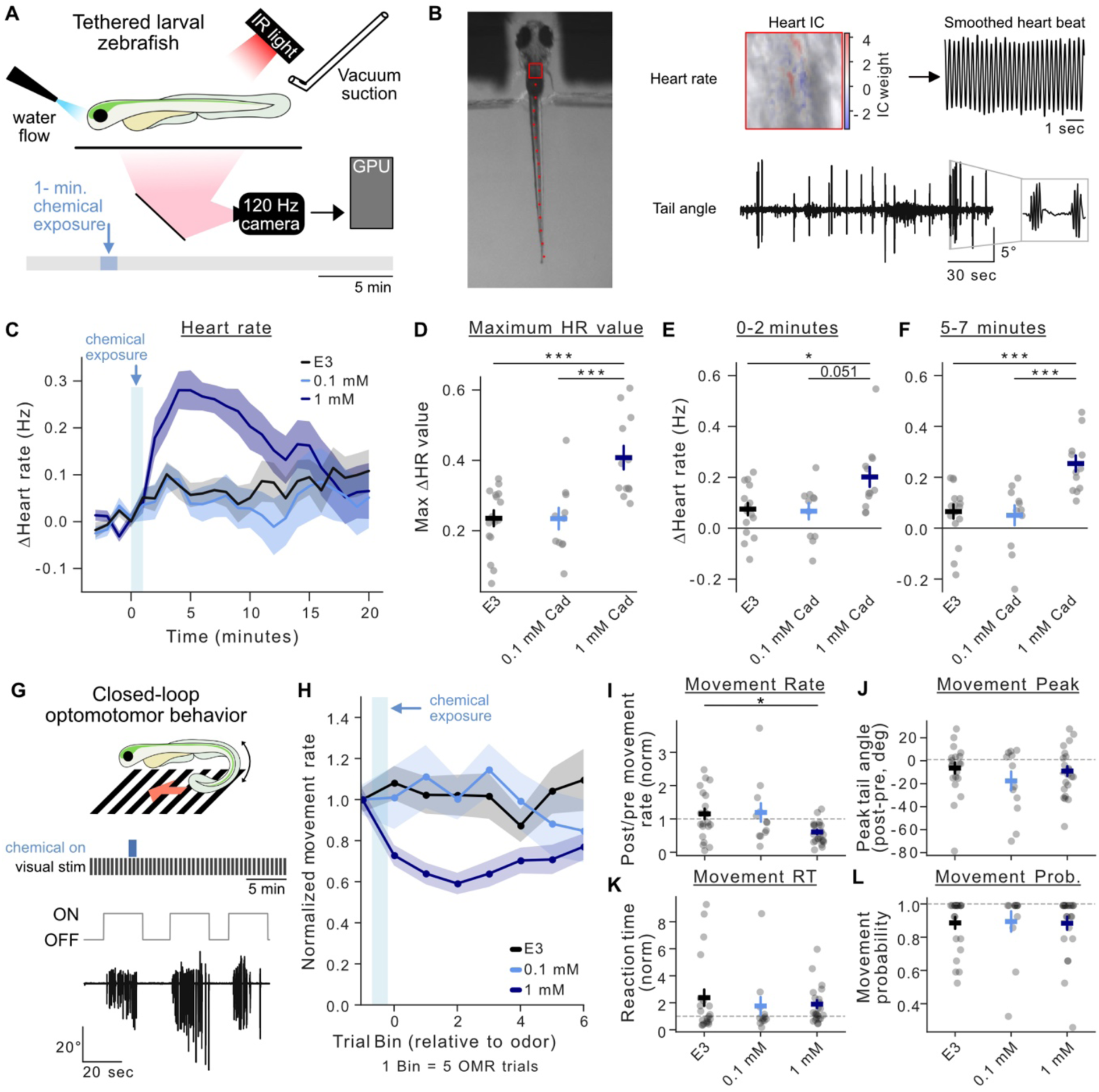
Decay-related chemicals induce a persistent state in larval zebrafish. (**A**) Behavioral paradigm of head-fixed odor presentation: odor delivery is controlled via pinch solenoid and removed by vacuum suction. (**B**) Image of larval zebrafish in the behavioral set-up showing the extraction pipeline for heart rate and tail movement. (**C**) Change in heart rate (Hz) over time of fish post-cadaverine onset. Change in heart rate binned over 1 minute (E3: n=16; 0.1 mM Cad n=11; 1 mM Cad n=12). (**D**) Maximum change in heart rate (Hz) post-odor onset from fish plotted in C. Error bars denote SEM,*** p<0.001. One-way ANOVA (F=11.920, p=0.0001) followed by Tukey’s multiple comparison *post hoc*. E3 vs 1mM Cad (p= 0.0002), 0.1 mM vs 1 mM Cad (p=0.0007). (**E**) Change in heart rate 0-2 minutes post-odor onset from fish plotted in C. Error bars denote SEM, * p<0.05. Kruskal-Wallis (H=8.357, p=0.0153) followed by Dunn’s test with Bonferroni correction. E3 vs 1 mM Cad (p=0.0251). (**F**) Change in heart rate 5-7 minutes post-odor onset from fish plotted in C. Error bars denote SEM, *** p<0.001. One-way ANOVA (F=11.627, p=0.0001) followed by Tukey’s multiple comparison *post hoc*. E3 vs 1 mM Cad (p=0.0004), 0.1 mM Cad vs 1 mM Cad (p=0.0005). (**G**) Schematic of optic flow behavioral paradigm to measure OMR. (**H**) Normalized movement rate over time post-Cad exposure. Each bin contains 5 OMR trials, which is approximately 2-2.5 minutes. (E3 n=22, 0.1 mM Cad n=12, 1 mM Cad n=24). (**I**) Normalized movement rate of 15 trials post odor-onset compared to 3 trials pre-odor. Error bars denote SEM,* p<0.05. Kruskal-Wallis (H=8.447, p=0.0146) followed by Dunn’s *post hoc* test with Holm correction. E3 vs 1mM Cad (p=0.0257). (**J-L**) Normalized movement parameters of 15 trials post odor onset compared to 3 trials pre-odor for peak change in tail angle (**J**), reaction time (**K**), and movement probability (**L**). Kruskal-Wallis tests identified no significance for these parameters across groups. Error bars denote SEM.

We measured the heart rate, eye movements, and tail movements from larvae exposed to control solution (E3 only), or cadaverine in E3 at 0.1 or 1 mM. While decaying fish can produce many fold-higher concentrations of cadaverine^23^, prior work has shown behavioral effects and olfactory bulb responses at concentrations of 1 mM or lower^23,33–37^. Compared with fish exposed to E3 and 0.1 mM cadaverine, fish exposed to 1 mM cadaverine displayed a significantly increased heart rate post-stimulus (**Figure 1C-D**). The average time to maximum baseline-normalized heart rate was 6.42±5.6 minutes (mean±sem, N=12 fish), after exposure to 1 mM cadaverine for one minute. We therefore analyzed changes in heart rate at 0-2 minutes or 5-7 minutes post-stimulus onset, to assess immediate and persistent changes in physiology, respectively. In these early and late epochs, fish exposed to 1 mM cadaverine increased their heart rate by 0.20±0.03Hz and 0.25±0.03Hz, respectively, compared to E3 (0.08±0.02Hz and 0.07±0.03Hz, respectively) and 0.1 mM (0.06±0.03Hz and 0.05±0.04Hz, respectively) (**Figure 1E-F**). This heart-rate increase persisted no longer than 15 minutes, and could be repeated by a second pulse of 1 mM cadaverine 20 minutes after the first exposure (in a different set of fish; **Figure S1C-D**), indicating that short pulses of this concentration of cadaverine did not affect overall fish health or subsequent physiological responses.

We next asked if this persistent cadaverine-driven increase in heart rate is found upon exposure to other chemical products of flesh decay and putrefaction, such as putrescine, which also elicits avoidance in other species^23,38,39^. Like cadaverine, putrescine elicited an increase in heart rate (0.20±0.03 Hz at 5-7 minutes post-exposure; **Figure S1E-G**). We wanted to understand if this persistent increase in heart rate is present to chemosensory threats^36,40^ unrelated to decomposition. Given that zebrafish inhabit fresh water, we tested how 1 minute exposure to salt water (50 mM sodium chloride in E3) affected the heart rate of tethered larvae. In agreement with prior work^41^, we observed that exposure to 50 mM salt caused an immediate increase in the larvae’s heart rate (0.24±0.04 Hz) during the first 2 minutes (**Figure S1E-F**). After this rapid-onset tachycardia, heart rate quickly returned to baseline – unlike the persistent effects of cadaverine and putrescine (**Figure S1G**). Therefore, brief exposure to high concentrations of the decay-associated odors cadaverine or putrescine has a persistent effect on larval zebrafish physiology by eliciting a sustained increase in heart rate.

Across all conditions, we did not observe significant changes to either spontaneous tail movements (**Figure S1H-J**) or eye movements (**Figure S1K-M**) during the recording. To determine if 1 mM cadaverine influences visually-evoked movements, we engaged tethered fish in a closed-loop optomotor response (OMR) behavior, where simulation of a backwards drift over bottom-projected visual gratings elicit compensatory forward swimming^42–45^ (**Figure 1G, S1N**). Fish were presented moving gratings in 20-second trials, with 10-15 second inter-trial intervals; we binned behavioral responses by trial, not time, to ensure tail responses to one trial were not split across multiple bins, then normalized movement to the bin immediately preceding onset of cadaverine (0, 0.1, or 1 mM) (**Figure 1H**). In the presence of the ongoing OMR, fish showed a sustained increase in heart rate to 1 mM cadaverine compared to E3 and 0.1 mM (**Figure S1P-R**). After 1-minute exposure to 1 mM cadaverine, fish showed a decrease in tail movement rate in the subsequent 15 trials (∼8 minutes) by 61±7% of baseline, whereas fish exposed to 0.1 mM cadaverine (119±28% of baseline) or E3 (115±16% of baseline) did not (**Figure 1I, S1O**). While the rate of tail movement was altered, we observed no significant differences in peak tail angle, reaction time, or movement probability between these groups (**Figure 1J-L**). Therefore, a one-minute exposure to 1 mM cadaverine evokes a persistent state, characterized by increased heart rate and suppressed visually-driven locomotion over ∼ 8 mins.

### Exposure to high-concentration cadaverine initiates brain-wide persistent activity

We next asked if this cadaverine-evoked persistent state is associated with patterns of neural activity on similarly long timescales. We performed volumetric brain-wide cellular-resolution 2-photon calcium imaging from *Tg(elavl3:H2B-GCaMP6s)* fish during and after 1-minute cadaverine exposure (**Figure 2A-C**), and registered each recording to the MapZeBrain atlas^46,47^ for anatomical segmentation (**Figure S2A-B**, see **Methods**). We recorded activity from 5420.4 ± 603.6 neurons per fish (mean ± sem, N=15 fish), exposed to either 0, 0.1, or 1 mM cadaverine (N=5 each). Fish did engage in the OMR in these experiments, and thus tail movements were spontaneous. To identify slow-timescale neural activity, we used spectral filtering to separate slow-timescale neural activity from each individual’s fast-timescale locomotor-associated activity (**Figures 2D, S2C,** see **Methods**). We observed widespread activation of neurons across brain regions, where the average peak response of neurons’ slow-timescale activity was significantly increased after exposure to 1 mM cadaverine, compared to 0.1 mM and E3. We observed this increase in neurons of the telencephalon, dorsal thalamus, tectum, superior medulla oblongata (MO), intermediate MO, and inferior MO (**Figure S2D**). To visualize the location of the most-active neurons, we mapped their location onto the reference brain. In fish exposed to high concentrations of cadaverine, we found the highest peak responses were apparent in the forebrain (largely within the telencephalon), as well as dorsolateral regions of the hindbrain (particularly the superior dorsal MO) (**Figure 2E**).

**Figure 2.**
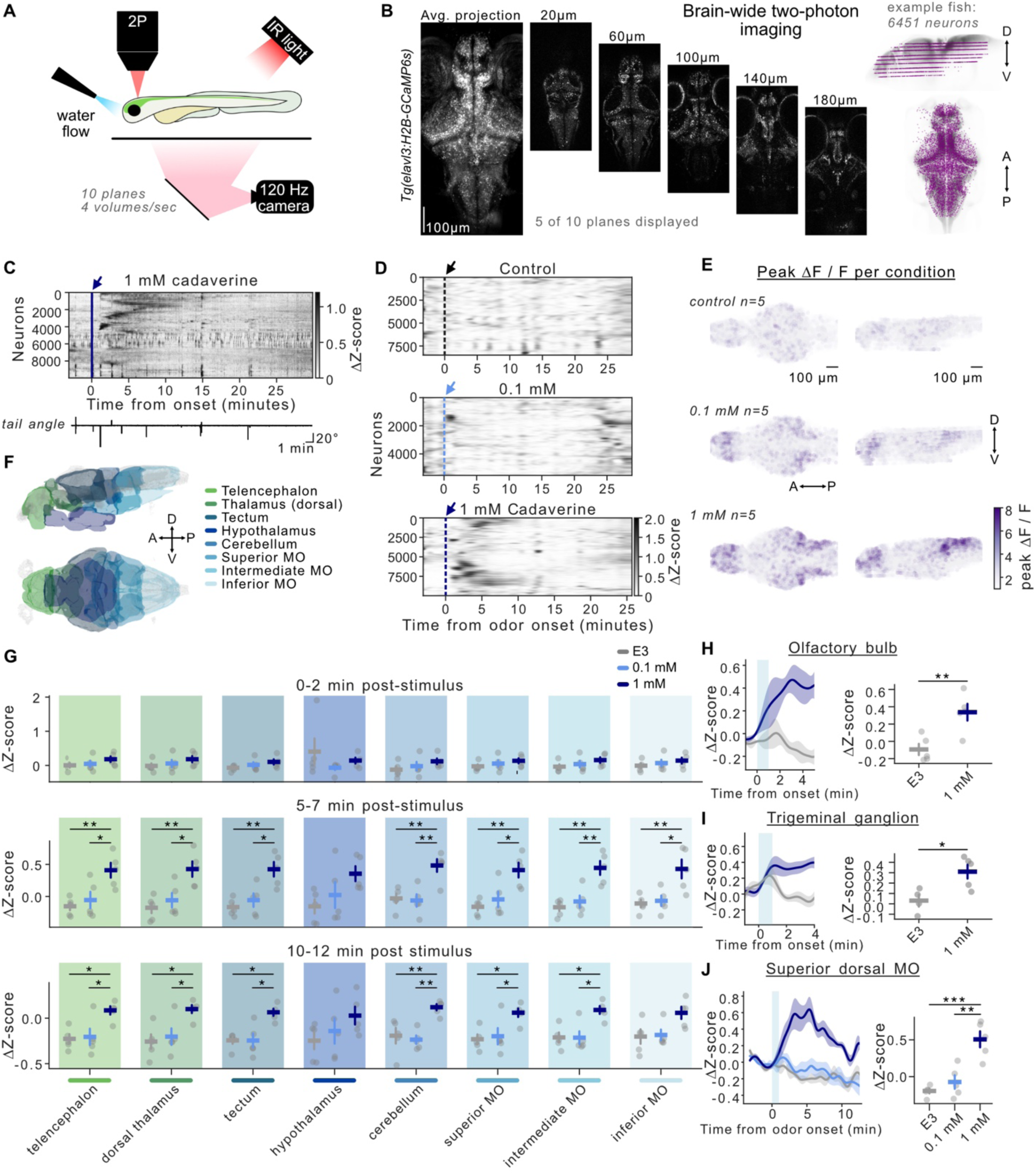
Brief cadaverine exposure initiates brain-wide persistent activity. (**A**) Brain-wide *in vivo* calcium imaging paradigm in head-fixed larval zebrafish. The setup is the same as Figure 1A, but moved under a two-photon microscope. (**B**) Sample imaging planes from brain-wide imaging volume showing fluorescent nuclear-localized calcium. Example dorsal and sagittal view of volumetric imaging to show location of planes (neuron locations in purple) in the brain. (**C**) Example Rastermap of neuronal activity from 1 fish exposed to 1 mM Cad with aligned tail angle trace (n= 9988 neurons). (**D**) Butterworth filtered slow frequency neurons from example fish for each condition (E3, 0.1 mM, and 1 mM Cad). (**E**) Peak change in fluorescence post-Cad exposure for all fish per condition, merged into a common reference brain. Signal intensity for each neuron is noted in the color scale at bottom right. (**F**) Illustration of broad anatomical brain regions spanning the larval zebrafish brain, listed anterior to posterior. (**G**) Average change in Z-score in a region per fish (n=5 per condition) in 0-2, 5-7, or 10-12 minutes post-stimulus. Error bars denote SEM, * p<0.05, ** p<0.01. Each condition was tested for normality (Shapiro-Wilk) and analyzed using a One-way ANOVA or Kruskal-Wallis test, followed by either Tukey’s or Dunn’s *post hoc* test with Holm correction for multiple comparison. 0-2 minutes: none. 5-7 minutes: telencephalon (ANOVA F=8.0160, p=0.0062; 1 mM vs E3 p=0.0073; 1 mM vs 0.1 mM p=0.0234), dorsal thalamus (ANOVA F =8.8646, p=0.0043; 1 mM vs E3 p=0.0049; 1 mM vs 0.1 mM p=0.0194), tectum (ANOVA F=10.6260, p=0.0022; 1mM vs E3 p=0.0026; 1 mM vs 0.1 mM p=0.0106), cerebellum (ANOVA F=11.1337, p=0.0018; 1 mM vs E3 p=0.0050, 1 mM vs 0.1 mM p=0.0032), superior MO (ANOVA F=8.8049, p=0.0044; 1 mM vs E3 p=0.0049; 1 mM vs 0.1 mM p=0.0206), intermediate MO (ANOVA F=13.7128, p=0.0008; 1 mM vs E3 p=0.0011; 1 mM vs 0.1 mM p=0.0036), inferior MO (ANOVA F=8.3696, p=0.0053; 1 mM vs E3 p=0.0083; 1 mM vs 0.1 mM p=0.0136). 10-12 minutes: telencephalon (ANOVA F=7.0552, p=0.0094; 1 mM vs E3 p=0.0144; 1 mM vs 0.1 mM p=0.0219), dorsal thalamus (ANOVA F=6.6177, p=0.0116; 1 mM vs E3 p=0.0142; 1 mM vs 0.1 mM p=0.0353), tectum (Kruskal-Wallis H=8.6400, p=0.0133; 1 mM vs E3 p=0.0327; 1 mM vs 0.1 mM p=0.0327), cerebellum (ANOVA F=10.4201, p=0.0024; 1 mM vs E3 p=0.0083, 1 mM vs 0.1 mM p=0.0034), superior MO (ANOVA F=5.7717, p=0.0175; 1 mM vs E3 p=0.0232; 1 mM vs 0.1 mM p=0.0429), intermediate MO (Kruskal-Wallis H=8.0600, p=0.0178; 1 mM vs E3 p=0.0473; 1 mM vs 0.1 mM p=0.0267). (**H**) Average trace of change in Z-score for 1 mM Cad and E3 in the olfactory bulb and the first 6 minutes post-stimulus onset. Shaded region denotes SEM, ** p<0.01. Welch’s t test (t = – 3.6682, p = 0.0080). (**I**) Average trace of change in Z-score for 1 mM Cad and E3 in the trigeminal ganglion and the first 6 minutes post-stimulus onset. Shaded region denotes SEM, * p<0.05. Welch’s t test (t =-3.1397, p=0.0164). (**J**) Change in Z-score for 1 mM, 0.1 mM, and E3 in the superior dorsal MO(ANOVA F=19.5125, p=0.0002, 1 mM vs E3 p=0.0011, 1 mM vs 0.1 mM p=0.0002.

We quantified the persistence of slow-timescale neural activity over three different epochs (0-2 minutes, 5-7 minutes, and 10-12 minutes post-stimulus) to correspond with the timescale of persistent changes in heart rate and OMR suppression (**Figure 2F-G**). During the first 2 minutes post-stimulus, 1 mM cadaverine showed no significant changes from E3 and/or 0.1 mM cadaverine in any brain region (**Figure 2G**, top), reflecting the sparse early activation of sensory neurons in these large anatomical regions. In the 5-7 minute epoch, fish exposed to 1 mM cadaverine had significant changes in z-scored activity spanning most regions of the brain, including the telencephalon, dorsal thalamus, tectum, cerebellum, and MO (**Figure 2G**, middle). Lastly, by 10-12 minutes post-stimulus, we continued to observe significant differences in 1 mM cadaverine between 0.1 mM and E3 for the regions listed above, excluding the inferior MO (**Figure 2G**, bottom). Additionally, across all regions and time periods examined here, slow-timescale neural activity in fish exposed to 1 mM cadaverine was higher than fish exposed to 0.1 mM cadaverine, with multiple hindbrain regions showing more sustained activity 5-7 minutes post-stimulus compared to more anterior regions (**Figure S2E**). In addition, we found that cadaverine exposure does not significantly affect the correlation of a neuron’s fast-timescale activity with spontaneous motor activity (**Figure S2F**). Together, these results indicate that brief cadaverine exposure causes a widespread increase in persistent neural activity.

While the activation we observed in the olfactory-associated forebrain is expected^23,33,48,49^ (including the olfactory bulb, **Figure 2H**), the engagement of the hindbrain suggests a potential role for alternative chemosensory pathways^32,50–52^. Notably, we found that 1 mM cadaverine activates neurons in the trigeminal ganglion, a bilateral population of neurons located posterior to the eyes, and ventral to most of the brain (**Figure 2I**). These neurons play an important role in chemosensation, including responding to noxious chemicals, CO2, and pH^51,53–55^. Using single cell tracings from MapZeBrain^46^, we visualized the morphology of neurons whose somas are inside the trigeminal ganglia. While few trigeminal neurons are present in this dataset, we observed one neuron’s neurites projecting to the area around the olfactory epithelium and face^56^, and multiple cells extending axons into the dorsolateral brainstem regions where we observed persistent activity after cadaverine exposure (**Figure S3A**). This suggests trigeminal neurons could be sensory cells and/or receive input from sensory neurons (including cranial nerve zero^53^), and project directly into the brainstem. We also confirmed cadaverine-evoked persistent activity in the sub-region of the hindbrain, the superior dorsal MO, where trigeminal neurons terminated (**Figure 2J**).

Therefore, trigeminal neurons possess a morphology that can directly link sensation in the face/nose with the hindbrain regions where we observed persistent cadaverine-evoked activity. While other sensory modalities could also be involved (for instance, chemosensory neurons in the mouth, gut, gills, or lateral line)^32,50–52,57^, the well-characterized dual role of trigeminal and olfactory chemosensation in vertebrates^30,31,51,58^ prompted us to focus on the relative roles of these two modalities in cadaverine-induced persistence.

### Cadaverine-evoked persistence does not require olfactory neurons

We tested if the cadaverine-induced persistent behavioral and neural effects we observed in larval zebrafish were mediated solely by olfaction, by ablating neurons in the olfactory epithelium that include those expressing the cadaverine-detecting receptor, TAAR13c. We used conditional metronidazole (MTZ)-nitroreductase (NTR) ablation to selectively ablate neurons expressing olfactory marker protein (*omp*), in *Tg(omp:gal4;UAS:NTR-mCherry)* larvae. While *omp* does not label all primary olfactory neurons in zebrafish, it does label the ciliated neurons expressing TAAR13c (ref. ^49^). We verified successful ablation of *omp* neurons using structural imaging of the olfactory epithelium, and functional imaging in downstream regions^46,59^. We found reduced mCherry expression in animals placed in 10 mM MTZ for 24 hours (**Figure 3A**), In addition, early cadaverine-evoked activity in the subpallium, the primary downstream target of olfactory bulb neurons^46,59^ (**Figure S3B**), was suppressed in fish with ablation of *omp+* neurons (**Figure 3B-D**).

**Figure 3.**
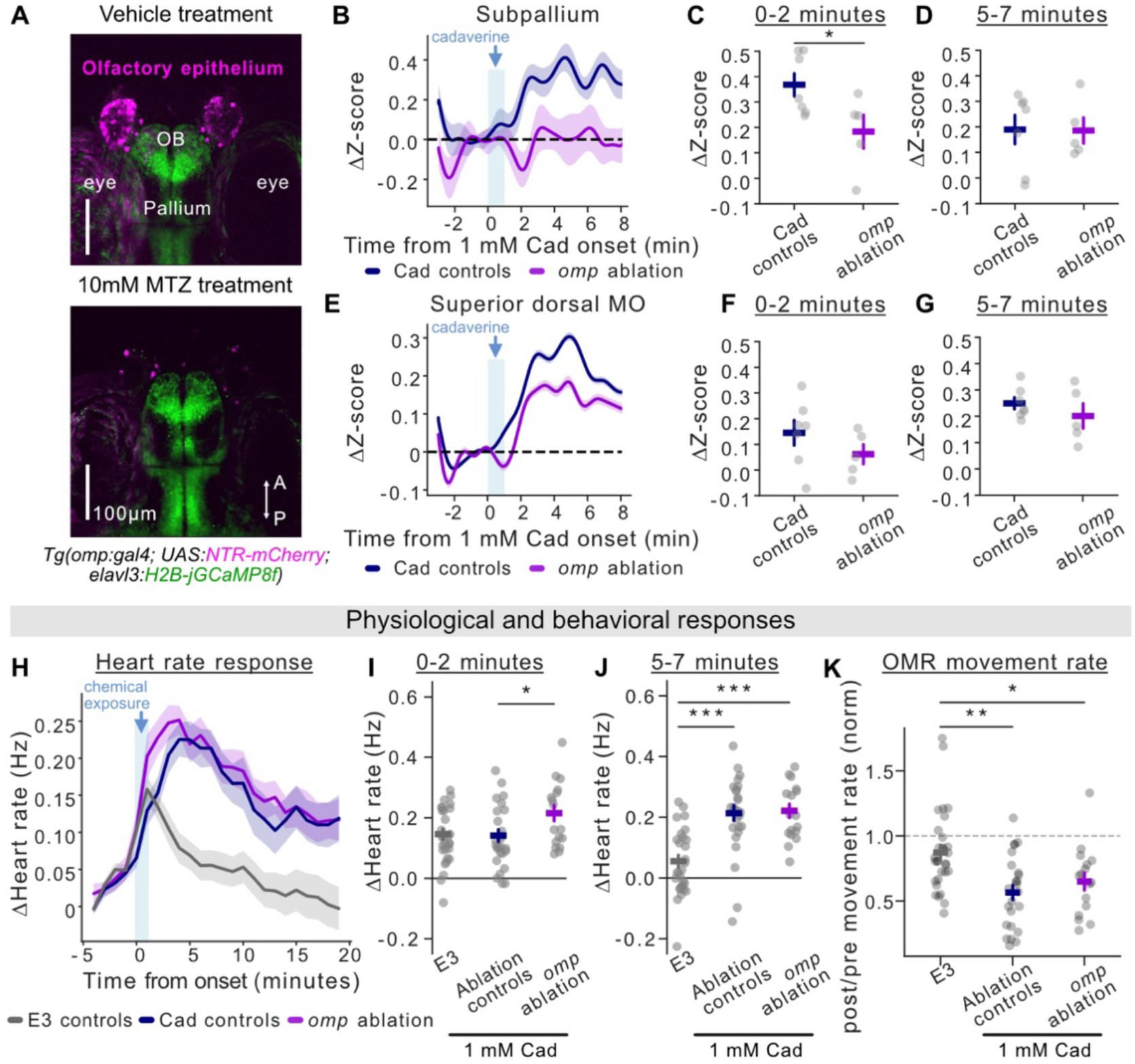
Cadaverine-evoked persistence does not require *omp*+ olfactory neurons. (**A**) Example image of *Tg(omp:gal4;UAS:NTR-mCherry;elavl3:H2B-GCaMP6s)* RFP+, GFP+ larvae treated with E3 or 10 mM MTZ. pan-neuronal jGCaMP8f (green); *omp+* neurons (magenta). (**B**) Combined fluorescence traces over time of fish in ablation control and *omp* ablated fish after exposure to 1 mM Cad in subpallium (n=6, 4). (**C**) Mean activity of non-motor related neurons in the subpallium 0-2 minutes after exposure to 1 mM Cad. Error bars denote SEM, *p<0.05 (Mann-Whitney U test, U=30.000, p=0.0480) (**D**) Mean activity of non-motor related neurons in the subpallium 5-7 minutes after exposure to 1 mM Cad. Error bars denote SEM, Welch’s T-test revealed no significant differences between groups. (**E**) Combined fluorescence traces over time of fish in ablation control and *omp* ablated fish after exposure to 1 mM Cad in superior dorsal MO (n=7,5). (**F**) Mean activity of non-motor related neurons in the superior dorsal MO 0-2 minutes post-Cad onset. Error bars denote SEM, Welch’s T-test revealed no significant differences between groups. (**G**) Mean activity of non-motor related neurons in the superior dorsal MO 5-7 minutes post-Cad onset. Error bars denote SEM, Welch’s T-test revealed no significant differences between groups. (**H**) Change in heart rate (Hz) over time post-stimulus (1 mM Cad or E3) onset. Change in heart rate binned over 1 minute. Shaded region denotes SEM. E3 controls combined treatment groups given a 1-minute pulse of E3 ([genotype, treatment]: RFP+, Vehicle; RFP-, MTZ, RFP+, MTZ; total n=29). Ablation controls combined treatment groups given a 1-minute pulse of 1mM Cad without *omp* ablation ([genotype, treatment]: RFP+, Vehicle; RFP-, MTZ; total n=26). *Omp* ablation consisted of RFP+ fish treated with MTZ; n=17). (**I**) Change in heart rate 0-2 minutes post-stimulus (1 mM or E3) onset. Error bars denote SEM, * p<0.05. One-way ANOVA (F=3.421, p=0.0383) followed by Tukey’s multiple comparison *post hoc* across same groups listed in H. Ablation controls vs *omp* ablation (p=0.0490). (**J**) Change in heart rate 5-7 minutes post-stimulus (1 mM or E3) onset. Error bars denote SEM, *** p<0.001. Kruskal-Wallis (H=26.163, p=2.08 x 10^-6^) followed by *post hoc* Dunn’s test with Holm correction. E3 vs Ablation controls (p=1.57 x 10^-5^); E3 vs *omp* ablation (p=0.0002). (**K**) Normalized movement rate of 15 trials post odor-onset compared to 3 trials pre-odor. Error bars denote SEM, * p<0.05; ** p<0.01. Kruskal-Wallis (H=13.138, p=0.0014) followed by Dunn’s *post hoc* test with Holm multiple comparison correction. E3 vs Ablation controls (p=0.0015); E3 vs *omp* ablation (p=0.0422).

After ablation or control manipulations (no NTR expression with MTZ incubation, or NTR expression without MTZ incubation), we conducted head-tethered experiments as described previously, and measured changes in heart rate, OMR behavior, and neural activity. Brain-wide imaging experiments revealed that cadaverine-evoked neural activity was still present after ablation of *omp+* neurons, with persistent activity in the superior dorsal MO (**Figures 3E-G, S3C**). In addition to our imaging results, we found no differences in persistent heart rate responses between ablation and control manipulations (**Figure 3H-J**). Furthermore, *omp*-ablated fish showed a significant suppression of the optomotor response post-cadaverine exposure (**Figure 3K**), similar to controls without ablations.

Therefore, while olfactory neurons of larval zebrafish respond to cadaverine (**Figure 2H**) and activate downstream targets in the forebrain^33–37,48^, we conclude that olfactory detection of cadaverine by *omp*+ neurons is not essential to induce the persistent effects of high-concentration cadaverine on the behavioral, physiological, or hindbrain activity measures we observe here. Consistent with our findings, a recent study reported that larval zebrafish showed no impairment in their avoidance of 1 mM cadaverine after laser ablation of olfactory neurons^33^. Therefore, these results indicate that 1 mM cadaverine is not solely detected by the olfactory system, but also impacts brain, body, and behavior through recruitment of other sensory systems, including trigeminal neurons (see **Figure 2I**).

### Chemosensory-induced persistence is mediated by the basic pH of concentrated cadaverine

The trigeminal ganglion of larval zebrafish is known to sense CO2, heat, pH, and noxious chemical irritants^52–54,60–62^. We hypothesized that pH may be a relevant factor, as previous work has reported a basic pH in small volumes of water with a decomposing fish^63,64^. Furthermore, we measured a pH of ∼9 for 1 mM cadaverine (and putrescine) in standard E3 solution, consistent with prior reports^33,34^, but a negligible effect of 0.1 mM cadaverine on the pH of E3 solution (∼7). To determine whether cadaverine’s basic pH can be reproduced with naturally decomposing zebrafish, we placed 30 dead zebrafish larvae into tubes with 1 mL of E3 and measured their pH over the course of 6 days (**Figures 4A, S4A-B**). We measured a basic pH in tubes with dead larvae within 24 hours, which continued to rise in the days afterward; this was not observed in tubes with E3 alone. Therefore, decomposition of fish postmortem increases the pH of the solution, at least within a small volume of water with limited buffering capacity.

**Figure 4.**
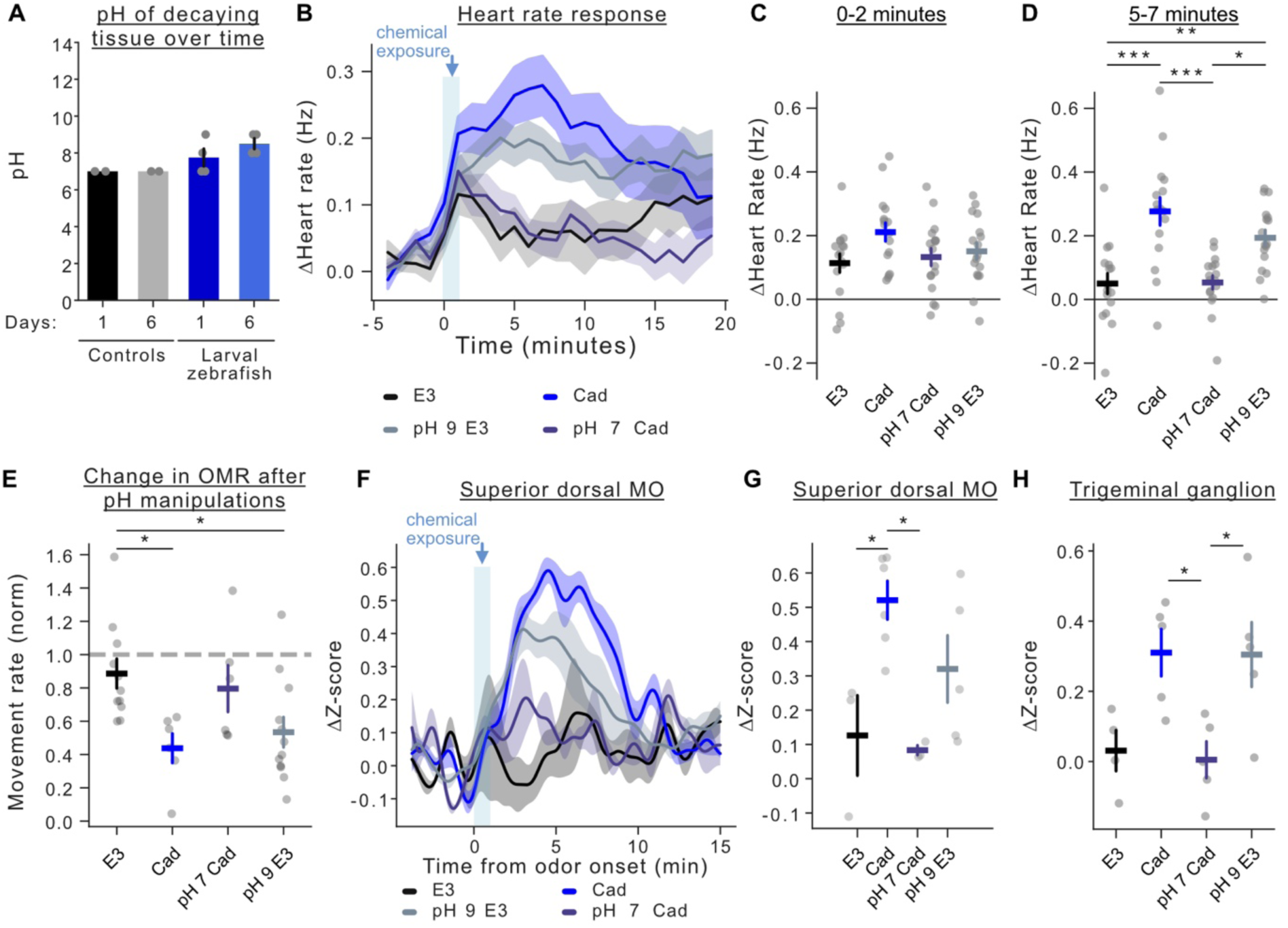
The persistent effects of high-concentration cadaverine are caused by its basic pH. (**A**) Blinded measurements of the pH of small water volumes for only E3 (Control) or with larval zebrafish on Day 1 and Day 6 post-mortem (**B**) Change in heart rate (Hz) over time post-stimulus onset. Change in heart rate binned over 1 minute. (E3 n=16; Cad n = 17; pH 7 Cad n=17, pH 9 E3 n=17). (**C**) Change in heart rate (Hz) 0-2 minutes post-stimulus onset from fish plotted in B. Error bars denote SEM. One-way ANOVA test identified no significance across groups. Error bars denote SEM. (**D**) Change in heart rate (Hz) 5-7 minutes post-stimulus onset from fish plotted in B. Error bars denote SEM, * p<0.05, ** p<0.01, *** p<0.001. One-way ANOVA (F=12.529, p=1.67 x 10^-6^) followed by Tukey’s multiple comparison *post hoc*. Cad vs pH 7 Cad (p=2.40 x 10^-5^); Cad vs E3 (p=2.37 x 10^-5^); Cad vs pH 9 E3 (p=0.0105); E3 vs pH 9 E3 (p=0.009) (**E**) Normalized movement rate of 15 trials post odor-onset compared to 3 trials pre-odor. Error bars denote SEM, * p<0.05. One-way ANOVA (F=4.286, p=0.0122) followed by Tukey’s multiple comparison *post hoc*. Cad vs E3 (p=0.0292); pH 9 E3 vs E3 (p=0.0397). (**F**). Normalized z-scored fluorescence of neurons in the superior dorsal MO.(**G**) Neural activity of superior dorsal MO to all conditions 5-7 minutes post-stimulus onset. Error bars represent SEM, * p<0.05. One-way ANOVA (F=6.107, p=0.0080) followed by Tukey’s multiple comparison *post hoc*. E3 vs Cad (p=0.0257); Cad vs pH 7 Cad (p=0.0133). (E3 n=3, Cad n=6, pH 7 Cad n=3, pH 9 E3 n=5). (**H**) Neural activity of trigeminal ganglion to all conditions (0-4 mins). Cad and E3 data previously shown in Figure 2I. Error bars represent SEM, * p<0.05. One-way ANOVA (F = 5.665, p = 0.0085), followed by Tukey’s multiple comparison *post hoc*. E3 vs Cad (p=0.07); Cad vs pH 7 Cad (p=0.0309); pH 9 E3 vs pH 7 Cad (p = 0.0344); E3 vs pH 9 E3 (p = 0.08). (E3 n=4, Cad n=5, pH 7 Cad n=5, pH 9 E3 n=5).

We therefore hypothesized that, at high concentrations, trigeminal neurons may be activated by the basic pH of cadaverine in solution. To disentangle the detection of a generalized high pH from cadaverine’s specific chemical properties and olfactory receptor binding, we used hydrochloric acid to lower the pH of cadaverine to match that of E3 (pH ∼ 7), or increased the pH of E3 with sodium hydroxide to match that of 1 mM cadaverine (pH ∼9). We then measured how heart rate, swimming in the OMR, and neural activity changed in response to these stimuli. In fish exposed to one-minute of pH 9 E3, we observed sustained elevation of the heart rate, on a similar timescale to that observed for 1 mM cadaverine (**Figure 4B-D**). Furthermore, when we neutralized 1 mM cadaverine to a pH of ∼7, we did not observe a significant change in heart rate compared with pH 7 E3 (**Figure 4B-D**). We therefore conclude that a basic pH is necessary and sufficient to elicit sustained heart rate elevation for 5-7 minutes after a 1-minute exposure to cadaverine. We also found that E3 raised to a pH of 9 caused a significant suppression of visually-driven swimming in the OMR, with a similar effect to fish exposed to 1 mM cadaverine. In contrast, when the pH of cadaverine is reduced to ∼7, we did not observe a suppression of the OMR (**Figure 4E**). These results indicate that brief sensation of water with a strongly basic pH is sufficient to suppress OMR over many minutes.

In addition to behavioral similarities between 1 mM cadaverine and pH 9 E3, we also observed similar persistent neural activity patterns between the two groups, which was not present upon exposure to pH 7 solutions (cadaverine or E3 controls; **Figure S4C-F**). We analyzed neural activity in the trigeminal-recipient superior dorsal MO (**Figure 4F**) and observed persistent activity in response to solutions of basic pH, but not neutral pH, regardless of the cadaverine concentration (**Figure 4G**). We also observed a similar pattern of stimulus-driven activity in neurons of the trigeminal ganglion (**Figure 4H**).

These results provide evidence that, in addition to canonical olfactory detection of cadaverine, an alternative chemosensory system detects the basic pH of high concentrations of cadaverine, or a high pH more broadly. We next explored potential cell-type specific sensory mechanisms mediating this high pH chemosensation.

### TrpA1b channels mediate high pH-evoked persistent behavioral effects

Similar to other vertebrates, zebrafish trigeminal neurons express polymodal cation channels, including TrpA1 and TrpV1^52,54,60^, which are reported to open in response to acidic or basic pH^65,66^. The TrpA1 paralog *trpa1b* is expressed in the trigeminal ganglion of larval zebrafish and is responsive to noxious chemical irritants such as the TrpA1 agonist allyl isothiocyanate, but does not influence thermosensory responses or hair cell function^54^. Furthermore, a prior study reported that exposing larval zebrafish to allyl isothiocyanate increases heart rate and anxiety-related thigmotaxis^67^. We therefore hypothesized that zebrafish sense cadaverine and other basic solutions via TrpA1b channels, to drive the persistent physiological and behavioral effects we have characterized here.

We used fluorescence *in situ* hybridization to label *trpa1b* mRNA in whole-mount larval zebrafish, and found *trpa1b* expression in the trigeminal ganglion (**Figure 5A**), consistent with previous studies^54^. Interestingly, we also saw *trpa1b* expression around the olfactory epithelium (**Figure S5A**), potentially corresponding to the soma of cells in cranial nerve zero^53^ that could respond to basic solutions or irritants presented to the front of the face. We hypothesized that cadaverine may activate the trigeminal ganglion via TrpA1b channels to drive changes in motor output and heart rate.

**Figure 5.**
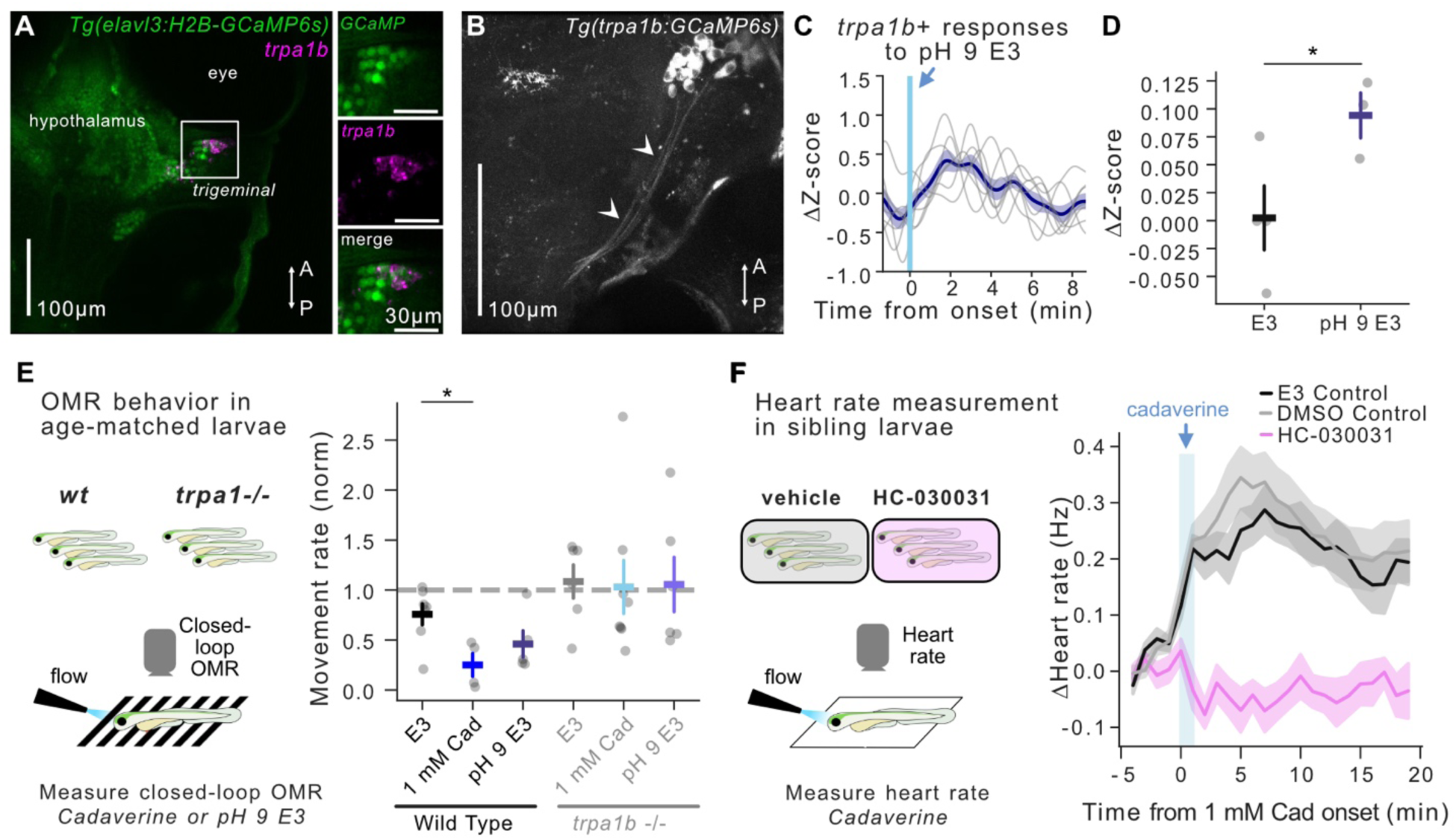
TrpA1b channels are required for cadaverine-evoked persistence. (**A**) *in situ* HCR of ***trpa1b+*** neurons (magenta) in the trigeminal ganglion of *Tg(elavl3:h2b-GCaMP6s)* fish (green). (**B**) Image of trigeminal ***trpa1b+*** neurons in *Tg(trpa1b:GCaMP6s)* fish. Arrowheads point to ***trpa1b+*** neuron axons descending towards the hindbrain (**C**) ***trpa1b+*** neuron responses from example fish exposed to pH 9 E3 for 1 minute. (**D**) Change in Z-score 0-6 minutes post-stimulus exposure (pH 9 E3, n=3; E3, n=4). Error bars represent SEM, * p<0.05. Welch’s T-test (t=-2.613, p=0.0484). (**E**) Normalized movement rate of wild type and knock out fish for 20 trials post odor-onset compared to 5 trials pre-odor. Error bars denote SEM, * p<0.05. One-way ANOVA (F=4.605, p=0.0308) followed by Tukey’s multiple comparison *post hoc*. Wild type: Cad vs E3 (p=0.0290); no significance detected between any trpa1b−/− fish conditions (Wild type: E3 n=9, Cad n=6, pH 9 E3 n=7; *trpa1b−/−:* E3 n=7, Cad n=8, pH 9 E3 n=7). (**F**) Change in heart rate (Hz) over time post-Cad onset. Change in heart rate binned over 1-minute, shaded region denotes SEM. (E3 Control n=13; DMSO Control n = 17; HC-030031 n=19).

We next conducted two photon imaging of the trigeminal ganglion in ***Tg(trpa1b:GCaMP6s)*** fish^62^ (**Figures 5B, S5B**), and found that trigeminal ***trpa1b+*** neurons respond to E3 solution at a basic pH (**Figure 5C-D**). To understand the role trpa1 channels play in OMR suppression, we measured the optomotor swimming behavior of larval zebrafish exposed to E3, 1 mM cadaverine, or pH 9 E3, and compared *trpa1b−/−* fish^62^ to age-matched wild type controls. While wild-type fish showed a persistent suppression of visually-driven swimming after 1 mM cadaverine, we found that *trpa1b−/−* fish did not change their OMR after chemosensory stimulation (**Figure 5E**). We therefore conclude that TrpA1b channels are required to drive the persistent suppression of OMR behavior after a 1-minute pulse of cadaverine.

Since ***trpa1b−/−*** fish were pigmented, we were unable to obtain direct optical measurements of heart rate. Instead, we presented 1 mM cadaverine and measured heart rate in unpigmented fish treated with either 10 µM of TrpA1b antagonist HC-030031 in DMSO (vehicle), E3 (E3 control), or vehicle alone (DMSO control) for 20 minutes before and during the behavior. While OMR behavior was more variable in drug-treated fish (**Figure S5C**), HC-030031 abolished cadaverine-induced heart rate elevation (**Figure 5F**).

In conclusion, we have found that at high concentrations, cadaverine is basic and activates a pH-sensing pathway via TrpA1b channels to initiate persistent behavioral and physiological changes. Our results implicate trigeminal chemosensation in this pathway, where *trpa1b*-expressing pH-responsive neurons project into the dorsolateral hindbrain (superior dorsal MO). We next sought to identify how this hindbrain input produces a persistent suppression of visually-driven swimming in the OMR.

### Chemosensory recruitment of persistent hindbrain inhibition suppresses movement

To examine how cadaverine exposure impacts visual processing and/or movement generation, we recorded neural activity while tethered fish engaged in the OMR task (**Figure S6A**). We performed brain-wide cellular-resolution calcium imaging in larval *Tg(elavl3:H2B-GCaMP6s)* zebrafish before and after 1 mM cadaverine exposure, and identified neurons whose activity was modulated by chemosensory stimulation, visual stimuli, and/or swimming actions (**see Methods, Figure S6B**). Visually-correlated neurons were largely seen in the tectum and pretectum, as has been previously reported^42–44,68,69^ (**Figure S6C**). We measured visually-responsive neurons across trials and observed a gradual decrease in visually-driven activity as the behavioral session progressed, but measured no difference between cadaverine– or E3-treated fish (**Figure S6D-E**). We therefore conclude that the suppression of visuomotor behavior in response to cadaverine is not due to inhibition of early visual processing in the tectum and pretectum.

Based on the persistent activity of motor-associated hindbrain regions (**Figure 2G**) and prior work demonstrating that hindbrain GABAergic neurons can inhibit visually-driven movement of larval zebrafish^10,45,70^, we hypothesized that some of the persistently-active populations of neurons we observe in the hindbrain may be inhibitory, and thus reduce swimming initiation or duration by inhibiting local neurons. We imaged the activity of GABAergic neurons in the hindbrain of *Tg(gad1b:Gal4; UAS:GCaMP6s)* zebrafish as they experienced one minute of E3, 0.1 mM, or 1 mM cadaverine (**Figure 6A**). We observed increased neural activity in *gad1b*+ neurons that persisted after exposure to 1 mM cadaverine (**Figures 6B-D, S6F-G**), suggesting these cells may serve to inhibit movement-generating neurons in the hindbrain on long timescales.

**Figure 6.**
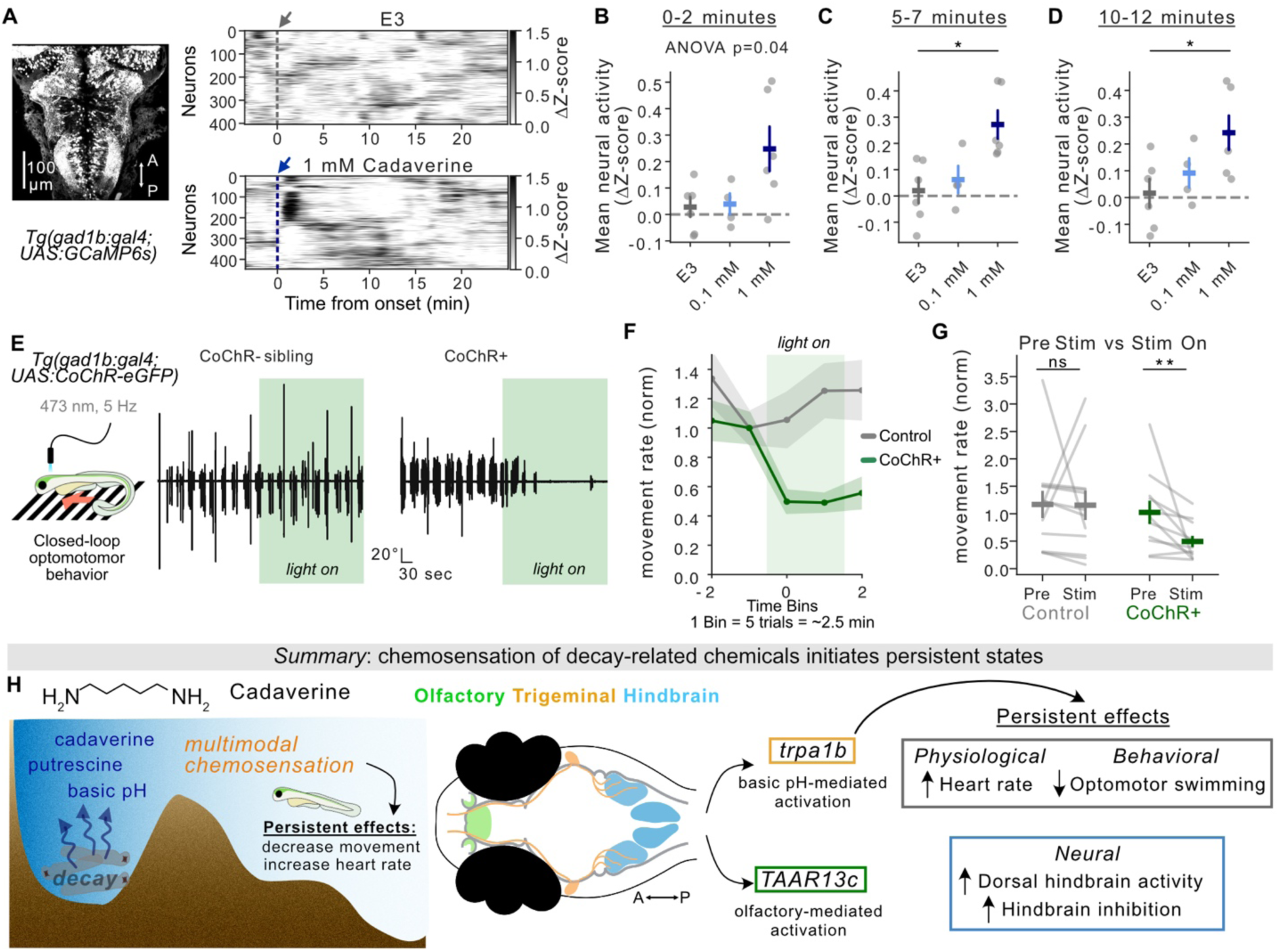
Chemosensory activation of persistent hindbrain inhibition suppresses movement. (**A**) Example Rastermaps of slow frequency GABAergic neuron fluorescence post-exposure to E3 or 1 mM Cad. (**B**) Mean fluorescence of GABAergic neurons 0-2 minutes post-exposure per condition (E3, 0.1 mM Cad., and 1 mM Cad.). Error bars denote SEM, One way ANOVA (F=4.142, p=0.0406) followed by Tukey’s *post hoc* for multiple comparisons. 1 mM Cad vs E3 (p=0.0513). (**C**) Mean fluorescence of GABAergic neurons 5-7 minutes post-exposure per condition. Error bars denote SEM, * p<0.05. Kruskal-Wallis (H=8.842, p=0.0120) followed by Dunn’s *post hoc* test with Holm correction for multiple comparison. 1 mM Cad vs E3 (p=0.013124). (**D**) Mean fluorescence of GABAergic neurons 10-12 minutes post-exposure per condition. Error bars denote SEM, * p<0.05. One way ANOVA (F=4.173, p=0.0398) followed by Tukey’s *post hoc* for multiple comparisons. 1 mM Cad vs E3 (p=0.0497); 0.1 mM Cad vs E3 (p=0.0314). (**E**) Experimental set up for optogenetic experiments with examples of tail traces for either CoChR+ or CoChR-siblings. (**F**) Normalized movement rate over time. Green shaded region means light was on to stimulate neurons. Each time bin contains 5 trials which is about 2.5 minutes. The shaded region surrounding the mean denotes SEM. (**G**) Normalized movement rate to pre-stim for pre-stimulation and during optogenetic stimulation of GABAergic neurons. Error bars denote SEM, ** p<0.01. Pre-stim contains Bins –2 and –1, Stim on contains bins 0 and 1. Wilcoxon test analyzed each group’s pre-stim vs stim-on movement rates; CoChR pre-stim vs stim-on (W=2.00, p=0.0029). (**H**) Summary schematic showing activation of **trpa1b+** neurons by decay chemicals and subsequent neural, physiological, and behavioral changes.

To test if the persistent activity of GABAergic neurons was sufficient to reduce movement in the absence of cadaverine or a basic-pH solution, we used optogenetics to activate hindbrain GABAergic neurons during optomotor behavior. We positioned an optic fiber over the hindbrain of transgenic *Tg(gad1b:Gal4; UAS:CoChR-eGFP)* fish (or opsin-negative controls) and pulsed a 473nm light at 5 Hz (**Figure 6E**, see **Methods**). As seen in prior studies^11^, we found that activation of hindbrain GABAergic neurons significantly reduced swimming to optomotor stimuli (**Figure 6F-G**). While activation of inhibitory neurons reduced movement rate in the OMR, we did not observe a change in heart rate (**Figure S6H**).

These findings suggest that the two features of this persistent state – locomotor suppression and tachycardia – may be implemented by distinct neural circuits that diverge after detection of cadaverine or basic water. This finding agrees with a recent study, reporting that while changes in movement and heart rate are negatively correlated, they are controlled by parallel pathways and can be manipulated independently^41^.

Together, these results demonstrate that the long-lasting suppression of optomotor swimming by brief cadaverine exposure is due, at least in part, to activation of persistently active GABAergic neurons in the hindbrain.

## Discussion

Animals possess a diverse array of sensory systems to detect and respond to salient features in their environment^1,2^, including cues indicating the presence of danger. Sensory detection of threats can elicit a persistent internal state that influences neural function and systemic physiology, allowing the animal to engage in adaptive behaviors on longer timescales, without the need for continual sensory input^4,5,7,12^. Here we have examined how the brief sensation of danger produces a persistent state, by studying the sensory and neural mechanisms of zebrafish larvae’s chemosensory detection of death and decay.

We found that a brief one-minute exposure to high concentrations of the decay-related chemical cadaverine evokes a long-lasting state in larval zebrafish characterized by an elevated heart rate and suppression of visually-driven locomotion over approximately ten minutes. Using brain-wide cellular-resolution calcium imaging, we found neurons whose cadaverine-evoked activity increased over the timescale of this persistent state, including cells within olfactory regions of the forebrain and a dorsolateral population of neurons in the hindbrain. While prior work has demonstrated that cadaverine is detected by TAARs in olfactory sensory neurons^23,33–37,48,49,71^, we found that ablation of olfactory neurons did not prevent the effects of high-concentration cadaverine on persistent hindbrain activity, heart rate, or locomotor suppression. Instead, we found these persistent responses are driven by the basic pH of cadaverine in solution^33,34^, rather than cadaverine itself. Similarly, decaying fish increase the pH of small water volumes^63,64^. We characterized this pH-sensing mechanism, and found that basic solutions activate neurons in the trigeminal ganglion, which project into the dorsolateral region of the hindbrain where we observed persistent activity. Trigeminal neurons express the polymodal cation channel TrpA1, and we found that genetic or pharmacological disruption of TrpA1 channels prevented cadaverine (or basic pH more generally) from inducing persistent suppression of the optomotor response. Furthermore, we find that the persistent optomotor suppression induced by the basic pH of cadaverine is driven by recruitment of local GABAergic neurons that inhibit visually-driven swimming. These results demonstrate that brief exposure to the chemical cues of death and decay can elicit a persistent state in larval zebrafish; we find that this is not driven by canonical olfaction, but by neurons in the trigeminal ganglia that sense the basic pH of decay-related chemicals at high concentration to produce a persistent internal state underlying altered behavior (**Figure 6H**).

While we have emphasized the role of the trigeminal ganglion here, our results do not preclude the possibility of contributions from other chemosensory systems. This includes taste^50^ and olfaction, where prior studies have demonstrated that cadaverine is able to elicit activation of the G-protein-coupled receptor TAAR13c at up to 1 mM^23,72^ and larval zebrafish will show avoidance of cadaverine solutions with a neutralized pH^35,37^. Therefore, while the specific olfactory detection of cadaverine and other decay-related polyamines may be sufficient to mediate avoidance (even at low concentrations^23,48^), here we propose that the persistent effects we observed (elevated heart rate, suppressed swimming, and hindbrain activation) are mediated by a general high pH-detection mechanisms via TrpA1 channels, including those expressed by trigeminal neurons. As decaying fish increase the pH of their surroundings in small water volumes^63,64^, olfactory and trigeminal systems may play different but complementary roles in responding to such threats: low-threshold olfactory detection of specific polyamines to drive an odor-specific short-timescale avoidance behavior, and a high-threshold trigeminal detection of non-specific basic pH to drive a generalized long-timescale state of alarm. Together, both chemosensory systems can contribute to adaptive behavior, with their relative contributions depending on ligand concentration, water buffering capacity, and other ecological factors^73,74^. The interplay between the trigeminal nerve and olfaction or taste has been previously explored across vertebrate species^26,30,31,51,75–77^, and further investigation of cadaverine detection will be aided by advances in methods for cell type-specific manipulation of receptor expression in zebrafish.

Here we reported that the persistent pH-driven effects of cadaverine are not present in fish lacking functional TrpA1b channels, and that *trpa1b*-expressing trigeminal neurons are responsive to cadaverine and high pH. However, we note that *trpa1b* is expressed in more cell types than just trigeminal neurons; these channels are present in the gut and lateral line hair cells^32,78^, and there is some evidence for *trpa1b* in cells around the olfactory epithelium^79,80^ (see **Figure S5A**). Therefore, multiple chemosensory populations may be responsive to high pH or other ligands that act on TrpA1 channels. Furthermore, the mechanisms of TrpA1 channel activation are not known. TrpA1 channels are diverse in structure and function^81^, and can respond to heat^82,83^, irritant and pungent chemicals^54,84–86^, light^82,87^, and other modalities^88^, depending on the cell types, species, and ecological context. Prior work has shown that polyamines such as cadaverine and putrescine differ from amino acids or bile salts in their actions on receptors^72,89^. In addition, TrpV1 and TrpA1 in cultured cells respond to both acidic and basic pH^66^, with the latter achieved via intracellular alkalinization^65,66^. Whether cadaverine’s effects are mediated by alkalization of TrpA1 at intracellular sites, other mechanisms of high pH detection^90^, and/or production of reactive electrophiles^91^, remains to be tested in future studies.

Finally, our results reveal a population of persistently-active neurons in the hindbrain that underlie this long-lasting state, but we have not determined the biophysical, neurochemical, and/or synaptic mechanisms that produce this persistence^12,92^. For instance, trigeminal neurons could release neuropeptides^93^ onto lateral hindbrain neurons to drive long-timescale intracellular metabotropic signaling^94–96^, recurrent synaptic excitation within the hindbrain could maintain long-timescale activity^4,9,97–99^, and/or glial cells could contribute to long-timescale modulation^10,70^. In addition, long-timescale feedback can include hormonal and synaptic interactions with other organs. A recent study found that spontaneous or salt-triggered increases in heart rate also result in suppression of optomotor swimming in larval zebrafish^41^, similar to our findings. However, optogenetic pacing of the larval zebrafish heart rate^100^ suggested that naturally-induced locomotor suppression and tachycardia are mediated by parallel circuits^41^. Future studies can examine the mechanisms of cadaverine-driven heart rate modulation, and identify how the brain integrates signals from across the body^101,102^ to trigger, inform, or maintain persistent states of the brain and body.

In conclusion, we have discovered and characterized a chemosensory pathway that detects brief exposures to high concentrations of cadaverine and basic pH, and drives persistent behavioral and physiological responses. This work highlights the multimodal nature of threat detection, where different sensory systems can be recruited to guide animal behavior depending on the severity of the threat and the context: at low concentrations of cadaverine fish use their olfactory system to avoid the stimulus, while at higher concentrations the pH increases recruit ***trpa1b+*** neurons in the trigeminal ganglion to initiate persistent behaviors. While the recruitment of multiple chemosensory modalities is by no means specific to larval zebrafish^30,31,75,76^, chemosensation in aquatic animals is multidimensional and subject to environmental variation in ion balance, pH, temperature, chemical composition, and oxygen/carbon dioxide ratios, among other factors. Given that many animals possess an array of diverse receptors to detect changes in chemical features, future work should account for stimuli that can impact multiple chemosensory systems to understand how sensory input shapes persistent behavior.

## Author contributions

GMP and ML-B designed experiments. GMP, MLW, MTN, and JJ conducted experiments. JSC and KAM contributed software and methods. GMP, MLW, and ML-B analyzed data. GMP and ML-B wrote the paper, with feedback from JSC and KAM.

## Acknowledgements

We thank members of the Lovett-Barron lab for feedback and support, Ajay Dhaka for *Tg(trpa1b:GCaMP6s)* and *trpa1b−/−* fish, Misha Ahrens for *Tg(elavl3:H2B-jGCaMP8f)* and *Tg(gad1b:Gal4)* fish, and Florian Engert for *Tg(omp:Gal4)* fish. We thank Lisa Stowers, Shrek Chalasani, and Brenda Bloodgood for advice, as well as Martin Tresguerres, Diana Rennison, and members of ADC-NiCE collaboration for feedback on this project.

## Funding

This project is supported by funds from the NSF GRFP (GMP and JSC), NIH T32GM133351 (MLW), the Allen Family Philanthropies through the Allen Discovery Center for Neurobiology in Changing Environments (ADC-NiCE; MLW and ML-B), the Helen Hay Whitney Foundation (KAM), as well as funds from the Searle Scholars Award, Packard Foundation Fellowship, Pew Biomedical Scholar Award, Klingenstein-Simons Fellowship in Neuroscience, the Sloan Research Fellowship, and NIH grants R00MH112840 and DP2EY036251 (ML-B).

## Statement on AI use

The authors used Claude (Anthropic), ChatGPT (OpenAI), and Gemini (Google) language models for coding assistance and consistency checks. The authors verified the output of models.

## Methods

| Reagent or Resource | Source | Identifier |
| --- | --- | --- |
| **Chemicals** |  |  |
| Cadaverine | Sigma Aldrich | CAS#462-94-2 |
| Putrescine | Fisher | CAS#110-60-1 |
| NaCl | Fisher | CAS#7647-14-5 |
| HC-030031 | Tocris | CAS#34085-38-7 |
| Metronidazole | Sigma Aldrich | CAS#443-48-1 |
| Hydrochloric acid | Fisher | CAS#7647-01-0 |
| Sodium hydroxide | Honeywell | CAS#1310-73-2 |
| **Experimental transgenic lines (*Danio rerio*)** |  |  |
| Tg(*elavl3:H2B-GCaMP6s*) | Freeman et al., 2014 (ref. <sup>103</sup>) | jf5Tg |
| Tg(*elavl3:H2B-jGCaMP8f*) | Zhang et al., 2023 (ref. <sup>104</sup>) | N/A |
| Tg(*omp:Gal4*);<br>Tg(*UAS:NTR-mCherry*) | Ferreira et al., 2014; Davison et al., 2007 (refs. <sup>105,106</sup>) | N/A |
| Tg(*gad1b:Gal4*);<br>Tg(*UAS:GCaMP6s*) | Forster et al., 2017; Chen et al., 2013 (refs. <sup>107,108</sup>) | zf3838Tg, mpn101Tg |
| Tg(*gad1b:Gal4*);<br>Tg(*UAS:CoChR-eGFP*) | Förster et al., 2017;<br>Antinucci et al.,<br>2020 (refs. <sup>107,109</sup>) | zf3838Tg, jf44Tg |
| *trpa1b*<sup>-/-</sup> | Lecamp et al., 2026<br>(ref. <sup>62</sup>) | N/A |
| Tg(*trpa1b:GCaMP6s*) | Lecamp et al., 2026<br>(ref. <sup>62</sup>) | N/A |
| Wildtype (AB) |  | ZIRC ZL1 |
| **Software and algorithms** |  |  |
| Bigstream | Wang et al.,<br>2021(refs. <sup>110,111</sup>) | https://github.com/JaneliaSciComp/bigstream |
| Suite2p | Pachitariu et al.,<br>2016 (ref. <sup>112</sup>) | https://github.com/Mouseland/suite2p |
| MapZeBrain | Kunst et al., 2019<br>(ref. <sup>46</sup>) | https://mapzebrain.org/home |
| Custom python code | This paper | Github upon publication |

**[utbl1]**

## Zebrafish

All procedures were approved by the University of California, San Diego’s Institutional Animal Care and Use Committee (IACUC). We used larval zebrafish for all experiments in this study, tested between 7 and 9 days post fertilization (d.p.f.). Animals were group housed in a 14-10-h light-dark cycle, temperature-controlled room and raised according to Zebrafish International Resource Center guidelines. All experiments were conducted during the light period (9:00-18:00). Larvae were fed rotifers twice daily from 5-14 d.p.f., and Artemia thereafter into adulthood. At this larval stage of development, the sex of zebrafish is not yet specified.

For all behavioral experiments and brain-wide neural imaging we used nuclear-localized, green calcium indicator: *Tg(elavl3:H2B-jGCaMP8f)* and *Tg(elavl3:H2B-GCaMP6s).* For GABAergic imaging we used: Cytosolic green calcium indicator: *Tg(gad1b:Gal4); Tg(UAS:GCaMP6s).* For channelrhodopsin expressed under *gad1b* promoter we used: *Tg(gad1b:Gal4); Tg(UAS:CoChR-eGFP).* For nitro-reductase expression under *omp* promoter we used a nuclear-localized, green calcium indicator: *Tg(omp:Gal4); Tg(UAS:NTR-mCherry); Tg(elavl3:H2B-jGCaMP8f).* For imaging *trpa1b*-expressing cells we used: *Tg(trpa1b:GCaMP6s).* For trpa1b knock-out behavioral experiments we used: *WT (AB) trpa1b−/−*.

## Behavioral experiments with and without optic flow

All behavioral experiments were controlled with BonsaiRx (bonsai-rx.org), using BonVision and BonZeb packages^113–115^ and custom routines to control and time stamp camera acquisition, odor delivery, visual stimuli, and/or optogenetic stimuli. Behavioral videos were acquired from a camera mounted below the fish, at 121 Hz (FLIR Grasshopper, #33-534) with a 12X Zoom lens (ThorLabs, MVL12X12Z) and long-pass filter (ThorLabs, FBH850-10). The fish was projected with infrared (IR) light (ThorLabs, M850L3), and bottom-projected white light (AnyBeam) for opto-stimulation and optomotor response (OMR) experiments or in the dark for simple odor delivery.

Zebrafish were mounted dorsal side up in a thin layer of 3% low-melting point agarose (Invitrogen) in the lid of a 35-mm Petri dish (Fisher) and positioned under a stereomicroscope (Leica, S9D). Once solidified, agarose was removed from around the tail posterior to the swim bladder and around the nose and mouth with a fine scalpel. Fish were then habituated in the Petri dish for 20 minutes in E3 solution (5 mM NaCl, 169 μM KCl, 330 μM CaCl2, 161 μM MgSO4·7H2O) at room temperature before behavioral testing. Afterwards, fish were moved to the behavioral apparatus and acclimated to a constant flow of E3 solution at 1.5mL min^-1^ heated to approximately 27°C with a heating apparatus (Automate Scientific, ThermoClamp-3) for 20 minutes. E3 solution flowed through a 200-um opening at the end of a pipette tip, positioned 20-30° in front of the fish and pointed at the nose and mouth. Flow switched between E3 solution, cadaverine (0.1, 0.5, or 1 mM added to E3), salt (50 mM added to E3), putrescine (1 mM added to E3) or blank (E3) using a pinch valve solenoid (Cole Palmer P/N 98302-02) controlled by a digital I/O device (National Instruments USB-6525). Behind the fish, a custom vacuum made from a bent borosilicate electrode (WPI) was placed at the top of the Petri dish to remove E3 and odor throughout the duration of the experiment. This configuration was used for standalone behavioral experiments and for experiments under the two-photon microscope. Each fish was exposed to 5 minutes of baseline, 1-minute of odor delivery, and 19 minutes of baseline.

To measure the optomotor response (OMR), closed-loop gratings were projected at 60 Hz to a diffuse screen under the fish in a trial structure. Each trial of the gratings was 20 seconds long with a 10-15 second inter-trial-interval. Trials were presented during the entire behavioral experiment, in a closed-loop paradigm^114^. Tail movements counteracted the backwards drift of the fish, by adding forward motion during swimming. This closed-loop paradigm encourages continuous visuomotor behavior^10,44^. For odor delivery experiments with OMR, the fish was recorded for 5 minutes of baseline, 1-minute odor delivery, and 19 minutes post-stimulus.

## Pharmacology experiments

For trpa1 antagonist experiments, mounted fish were placed in 10µM HC-030031, 0.1% DMSO for 20 minutes during acclimation. OMR experiments were conducted with HC-030031 in the water flow. Fish were recorded for 5 minutes of baseline, 1-minute odor delivery, and 19-minutes post-stimulus presentation.

## Optogenetic experiments

For CoChR stimulation experiments, fish were embedded as described above and exposed to 1.5mL min^-1^ flow of E3 solution. Neurons were stimulated with a 470-nm fiber coupled LED (ThorLabs, M470F3) through a 100-µm 0.66NA fiber (Doric Lenses) positioned above the hindbrain at 45-60° with a micromanipulator (ThorLabs, DT12). The optic fiber made slight contact with the agarose encasing the fish. Each fish was exposed to 3 stimulation trials. Light was delivered as 100-ms pulses at 5 Hz for 5 minutes (5-minute inter-trial interval).

## Chemogenetic neuron ablation

Tg(*omp:Gal4*) fish were sorted at 5 d.p.f. for NTR-mCherry expression using an upright fluorescence dissecting microscope (Leica, MZ10F). mCherry+ fish and an equal number of pigment-matched mCherry-clutch mate controls were collected and placed into 10cm Petri dishes with E3 containing 10mM metronidazole. Dishes were covered in tin foil and removed 24 hours later at 6 d.p.f. Fish were then inspected to ensure decreased or absent mCherry fluorescence before continuing the experiment. mCherry fluorescence was also checked on testing day. Fish were transferred to a new 10cm Petri dish with standard E3, provided rotifers, and allowed to recover until testing at 7, 8, or 9 d.p.f.

## pH measurements of decaying fish

7 d.p.f. larval zebrafish were separated into 1.5mL tubes (∼30 fish per tube) and sacrificed on ice for 30 minutes. Tubes were filled with 1mL of either E3 left for 24 hours before measuring the pH of the solution. pH was measured again on day 6. Matching of pH strip color was conducted by an observer blinded to the condition.

## Behavioral analysis

We analyzed time series of multiple variables collected at a common 121 fps sampling rate (max of camera) from Bonsai recordings: time, odor control solenoid voltage, visual gratings movement, angle between the top and bottom extent of the tail (detected online using BonZeb^114^), and video of a cropped section around the eyes and heart.

In each frame of the video, frames were cropped to include just the eyes using openCV, downsampled to 90×60 pixels, and thresholded to identify two dark shapes in each frame (cv2.findContours). These shapes were fitted with an ellipse (cv2.fitEllipse), and the angle of each ellipse’s major axis was saved (one measurement per video frame). The time series of each eye’s angles were smoothed by a Hanning filter (120 frame window). We then identified the peaks of eye movements (left or right) that exceeded a threshold of 1.5 standard deviations above the mean, and used these movement peaks to calculate a saccade rate. To compare across subjects, we then binned saccade rates in 1-minute intervals, with the mean rate value in these bins subtracted by the mean of the 5 mins preceding the odor onset.

To quantify heart rate, we first used openCV to crop the image of the heart from the video, and downsample to 30×30 pixels. We then flattened this array in each video frame, and performed an independent components analysis (sklearn.decomposition.FastICA, n_components=5 or 10). We then manually selected the component with the strongest oscillatory signal over the entire video (at the approximate heart rate of larval zebrafish). Often, multiple components had oscillatory signals, and were offset in phase (i.e.: for movement of the 2 heart chambers). The selected independent component signal was smoothed with a Hanning filter (22 frame window), normalized to 0-1, and peaks were detected using scipy.signal.find_peaks (height=0.8, distance = 150 ms). Heart rate was then calculated in 1s bins, and smoothed in a sliding window of 5s over this array of beat times. To compare across subjects, we then binned heart rates in 1-minute intervals, with the mean rate value in these bins subtracted by the mean of the 5 mins preceding the odor onset.

Movement bouts were calculated from tail angle recordings, by classifying segments of time where the absolute tail angle exceeded a threshold of 1.5 standard deviations above the mean. Adjacent bouts were separated from each other if there was 500ms of tail movement below the threshold. Bouts could range from 25 ms to 10s. To compare across subjects, we then binned movement rates (i.e.: bout onset rates) in 1-minute intervals, with the mean rate value in these bins divided by the mean of the 5 mins preceding the odor onset.

Fish were excluded from further analysis if we failed to obtain measurements of heart beats (for instance, if the heart was obscured and no reliable oscillatory signal was present in the top 10 independent components), or tail movements (for instance if the tail was not properly isolated during data collection).

In experiments with optomotor stimuli, we indexed the onset of each trial so that trial 0 is the first OMR trial after odor onset. For each trial, we quantified the mean movement rate in the 18s after stimulus onset. We quantified the visually-driven movement rate in trials 1-16 after the odor onset (excluding trial 0), divided by the mean movement rate in the three trials preceding odor onset. We used these parameters for all experiments, except for the experiments in knockout fish, where we used a longer pre-stimulus period (5 trials preceding odor onset) and post-stimulus period (trials 2-20 after odor onset). We only analyzed fish where the pre-odor movement rate was at least 1 Hz.

## Two-photon calcium imaging

Two-photon Ca^2+^ imaging was performed with a ThorLabs Bergamo II multiphoton microscope with a resonant scanner in bidirectional scanning mode^116^, controlled by ThorImageLS 4.3 and illuminated by an fs-pulsed 80-MHz Ti:S laser (MaiTai DeepSee; Spectra Physics). For brain-wide live imaging experiments at 930nm and ≤10mW excitation power, GCaMP8f and GCaMP6s fluorescence was imaged with a 16×/0.8W Nikon objective at x1 zoom in 10 *z*-planes, cropped to 800 x 384 pixels covering a field-of-view of 833 x 400µm, separated by 20µm z-spacing, at 3.6 volumes/s. Prior to behavior and functional brain imaging, a structural stack was obtained at 860nm or 930nm, with 1µm spacing and 25x frame averaging, starting 20µm above the first *z*-plane, ending 20µm below the last *z*-plane.

For hindbrain cytosolic GCaMP6s imaging, we used a 16x/0.8W Nikon objective at 1.1x zoom in 6 *z*-planes, cropped to 384×384 pixels (422×422µm), separated by 10µm, at 5.67Hz. Prior to behavior and functional brain imaging, a structural stack was obtained at 860nm or 930nm, with 1µm spacing and 25x frame averaging, starting 10µm above the first *z*-plane, ending 10µm below the last *z*-plane.

For imaging experiments of the trigeminal ganglion, we used a 16x/0.8W Nikon objective at 4x zoom with 6 z-planes, cropped to 512×512, separated by 10µm, at 2.93Hz. Prior to behavior and functional brain imaging, a structural stack was obtained at 930nm, with 1µm spacing and 25x frame averaging, starting 20µm above the first *z*-plane, ending 20µm below the last *z*-plane. Neural data was collected on either the left or right side.

Imaging experiments were conducted together with behavioral measurements, using the same head-tethered arena as described above.

## Fluorescence *in situ* hybridization

Probes for *trpa1b* were designed from previously published sequences^54,62^. Methods for *in situ* hybridization are adapted from prior studies^47^. Briefly, we designed hybridization probes using the split-initiation approach of *in situ* hybridization chain reaction HCRv.3.0.^117^, where 25-nucleotide DNA oligonucleotide pairs with split B1 initiation sequences were tiled across the length of the mRNA transcript, synthesized by Integrated DNA Technologies. De-conjugated hairpins (B1-647) were purchased from Molecular Technologies. Zebrafish were fixed overnight in 4% paraformaldehyde in 1xPBST at 4°C. After washing 3x in 1xPBST, larvae were permeabilized in 100% (v/v) methanol at –20°C for 10 minutes then rehydrated in 1:1 100% (v/v) methanol, 2x SSCT, then twice with 2xSSCT; 5 minutes each. Hybridization with split probes was done overnight at 37°C in 2xSSCT, 10% (v/v) dextran sulfate, 10%(v/v) formamide at a probe concentration of 4 nM. The next day, larvae were washed at 37°C in 2xSSCT, 30% (v/v) formamide then 2 times in 2xSSCT at room temperature, each for 20 minutes. During this time, dye-conjugated hairpins were heated to 95°C for 1 minute then snap-cooled to room temperature. Samples were placed in the 50µL of amplification buffer with B1 probe concentrations at 120nM overnight in the dark. Larvae were washed 3 times with 2xSSCT for 20 minutes then mounted in 3% low-melting-point agarose, covered in PBS or E3, and imaged under the two-photon microscope. Fish were imaged at 860nm and 1200nm in unidirectional resonant scanning mode (25x line average).

## Two-photon calcium imaging data processing

We only proceeded with analyses where fish remained healthy at the end of recording and did not have substantial z-movements. Imaging data were motion corrected in Suite2p, followed by cell extraction^112,118^. Time series were inspected^119^ to ensure there was no z-motion contamination or slow drift, and only included neurons with continuous measurements and which were classified as a cell by Suite2p (“iscell”=1). Neuron locations were aligned to a common brain atlas, as described below.

Behavioral time series (analyzed as described above) were cropped to the onset and offset of the 2-photon data acquisition periods. We excluded the first 5% of behavioral and neural measurements in each imaging experiment to avoid the influence of sound-evoked activity upon the initiation of scanning. Stimulus timing and calcium traces were aligned and resampled to a common 5 Hz sampling rate for further analysis. Each neuron’s fluorescence trace was smoothed with a 3-sec rolling mean and z-scored (scipy.stats.zscore). For the spatial maps in **Figure 2E**, we plotted neurons as peak F-F0/F0, where F0 is the mean of the trace before the odor onset.

To separate fast movement-related activity from slow “state”-related activity in each trace, we applied a low-pass Butterworth filter (scipy.signal.butter, N=4, slow_cutoff=0.01s, low-pass) or a high-pass filter (scipy.signal.butter, N=4, fast_cutoff=2.0s). From fast traces, we quantified each neuron’s Pearson correlation coefficient to the movement onset array (convolved with an exponential kernel with a 3.0s decay). For slow traces, we quantified the mean activity as 0-2 mins after odor onset, 5-7 mins after odor onset, and 10-12 mins after odor onset, and/or a sole 0-6 mins epoch.

For analysis of neural activity during visuomotor behavior, we used two approaches. To visualize task-responsive neurons, we fit a linear model (sklearn.linear_model.ElasticNetCV, cv=3, alphas=np.logspace( –3, 1, 5), l1_ratio=np.arange(0.1, 1.01, 0.25), selection=’cyclic’, max_iter=1000) to each neuron’s fluorescence time series. Regressors were visual stimulus times, visual stimulus onset times, odor times, odor onset times, and an array of movement onset times – all convolved with an exponential kernel (3.0s decay). We then plotted all neurons and colored them by the coefficient for each regressor. To compare visually-driven activity before and after odor, we measured the change in z-scored activity from the 5s before visual stimulus onset to the 20s after. We then divided the mean of this value after the odor (25 trials) by the mean before the odor (5 trials).

## Volume registration

Recorded neurons were registered to a common brain atlas for anatomical identification (MapZeBrain)^46,47^. The average of each functional z-plane’s motion-corrected time series was first assigned to a z-position within that fish’s high-resolution anatomical stack (see above), assigned by finding the maximum of the cross-correlation between planes. This assigned a z-value to the x-y coordinates of extracted neurons within that plane. The anatomical stack was aligned to the MapZeBrain standard brain (nls-GCaMP), using rigid and non-rigid registration in BigStream^110,111^. After each brain had reached sufficient alignment with the atlas, we applied these transformations to the x/y/z positions of recorded neurons.

Parameters are:

> *affine_kwargs = {‘initial_condition’: ‘CENTER’, ‘alignment_spacing’:2.0, ‘shrink_factors’:[32,16,8,4,2], ‘smooth_sigmas’:[32,16,8,4,2], ‘optimizer_args’:{’learningRate’:1.5,’minStep’:0., ‘numberOfIterations’:100}} deform_kwargs = {’control_point_spacing’: 256, ‘alignment_spacing’: 2.0, ‘control_point_levels’:(16,8,4), ‘shrink_factors’:[8,4,2], ‘smooth_sigmas’:[8,4,2], ‘optimizer_args’:{’learningRate’:0.8,’minStep’:0.,’numberOfIterations’:20,}}*

We performed this procedure for all fish recorded during brain-wide imaging. We applied this same procedure to brainstem recordings in *Tg(gad1b:Gal4;UAS:GCaMP6s)* fish, but changed the reference brain to “mpn155Tg”, and applied a mask (x = 450:900, y = 100:500, z = 0:280) to only register to the brainstem region. Trigeminal recordings were not registered to an atlas. Instead, cells in the trigeminal ganglion were manually identified (based on anatomical characteristics in *Tg(elavl3:H2B-GCaMP6s)* fish and location of neurons in *Tg(trpa1b:GCaMP6s)* fish) and selected for further analysis.

In brain-wide imaging experiments, we excluded any cells whose location fell outside the brain, as defined by the MapZeBrain regions “prosencephalon” (excluding “eyes”), “mesencephalon”, and “rhombencephalon”. We then classified cells as overlapping with the following anatomical regions: ‘telencephalon’, ‘thalamus_(dorsal_thalamus)’, ‘tectum’, ‘hypothalamus’, ‘cerebellum’, ‘superior_medulla_oblongata’, ‘intermediate_medulla_oblongata’, ‘inferior_medulla_oblongata’.

We also used the MapZeBrain atlas to download and plot neurons from the single-cell labeling dataset^46^, using neurons whose cell bodies reside in the olfactory bulb or the trigeminal ganglia.

## Statistics

Groups were tested for normality using the Shapiro-Wilk test. Non-parametric tests (Mann-Whitney U, Kruskal-Wallis H) were conducted if the Shapiro-Wilk p <0.05 for any group. Otherwise, parametric tests were used. *Post hoc* tests were conducted if the main effect was p<0.05. All *post hoc* tests (Dunn’s test for nonparametric data, or t-tests for parametric data) were corrected for multiple comparisons with a Holm correction. Statistical tests on neural activity measurements are averages at the level of individual subjects. Exact tests and p values are reported in the figure legends.

## Supplementary Figures

**Supplemental Figure 1.**
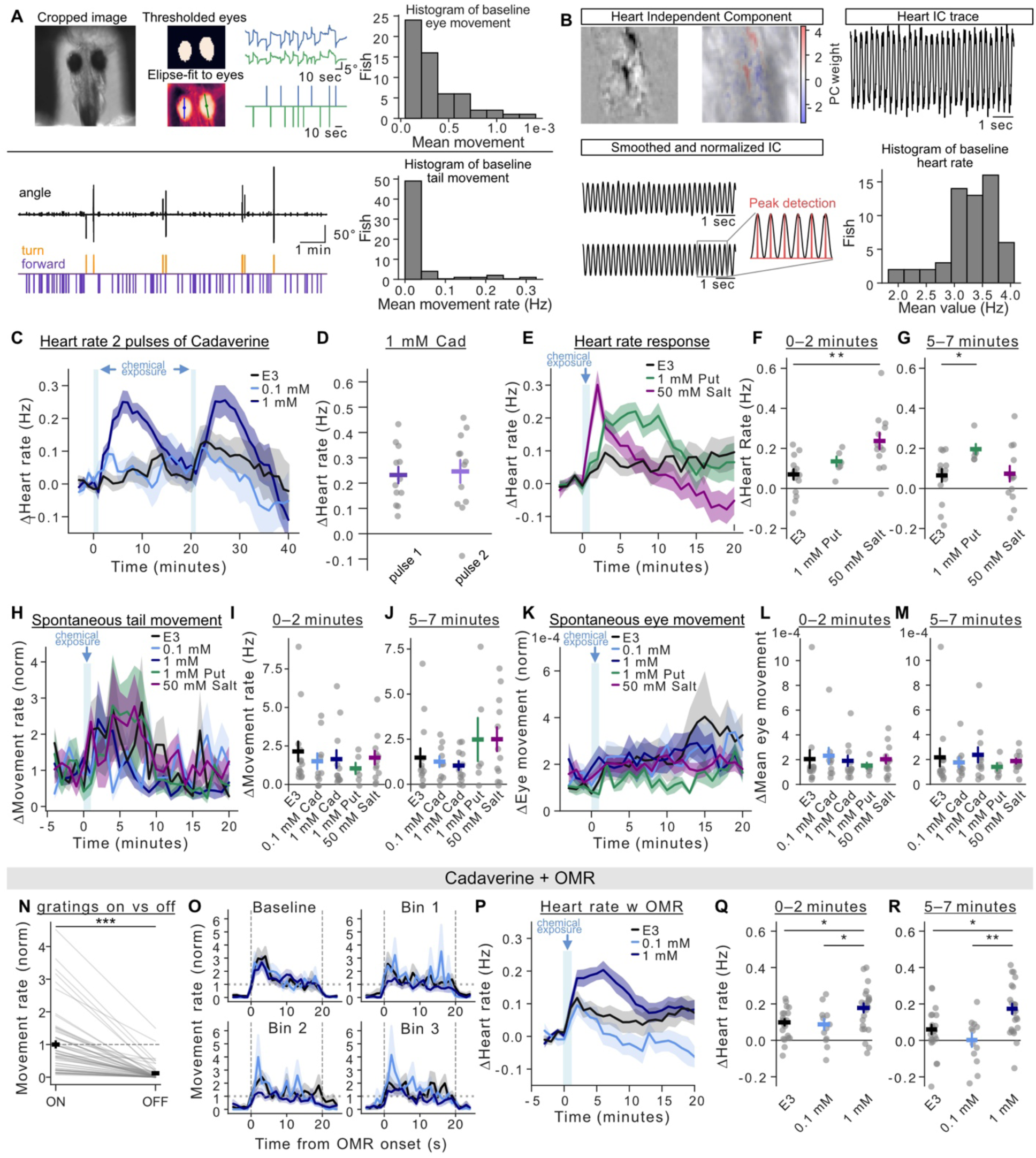
(**A**) Eye saccade detection and tail movement classification pipeline with histograms of baseline means across all fish (n=57). (**B**) Heart rate extraction pipeline with an independent component of heart rate pulled out, smoothed, and normalized for peak detection. Histogram shows the baseline heart rate across all fish (n=57). (**C**) Normalized heart rate traces to two pulses of the same cadaverine concentration (E3 n=13, 0.1 mM Cad n=11, 1 mM Cad n=12). Each fish heart rate is binned to 1 minute. (**D**) Change in heart rate to 1 mM Cad 5-7 minutes after each pulse. Error bars denote SEM. A paired t-test identified no significant difference in heart rate after each pulse. (**E**) Normalized heart rate responses to 1 mM Put (n=6) and 50 mM Salt (n=13). (**F**) Change in heart rate 0-2 minutes post-stimulus. Error bars denote SEM, ** p<0.01. Welch’s t-test: E3 and 50 mM Salt (T=-3.469, p=0.0023). (**G**) Change in heart rate 5-7 minutes post-stimulus. Error bars denote SEM, * p<0.05. Mann-Whitney U: E3 and 1 mM Put (U=15.000, p=0.0256). (**H**) Normalized spontaneous movement rate across all conditions. (**I**) Change in movement rate (Hz) 0-2 minutes post-stimulus. Error bars denote SEM. Kruskal-Wallis tests identified no significance for these parameters across groups. (**J**) Change in movement rate (Hz) 5-7 minutes post-stimulus. Error bars denote SEM. Kruskal-Wallis tests identified no significance for these parameters across groups. (**K**) Normalized spontaneous eye movement across all conditions. (**L**) Change in eye saccades 0-2 minutes post-stimulus. Error bars denote SEM. Kruskal-Wallis tests identified no significance for these parameters across groups. (**M**) Change in eye saccades 5-7 minutes post-stimulus. Error bars denote SEM. Kruskal-Wallis tests identified no significance for these parameters across groups. **(N**) Normalized change in movement when projected gratings are on (closed-loop, backwards drifting) or off (no gratings present). Taken from 5-trials pre-odor onset across all fish (n=57). (**O**) Average normalized movement rate over time per bin for all conditions. (**P**) Normalized heart rate traces to cadaverine concentrations while engaging in OMR (E3 n=22, 0.1 mM Cad n=12, 1 mM Cad n=24). (**Q**) Change in heart rate 0-2 minutes post-odor onset from fish plotted in P. Error bars denote SEM, * p<0.05. One-way ANOVA (F=4.450, p=0.0162) followed by Tukey’s multiple comparison *post hoc* test. E3 vs 1 mM Cad (p=0.0350); 0.1 mM vs 1 mM Cad (p=0.0477) (**R**) Change in heart rate 5-7 minutes post-odor onset from fish plotted in P. Error bars denote SEM, * p<0.05, ** p<0.01. One-way ANOVA (F=7.989, p=0.0009) followed by Tukey’s multiple comparison *post hoc* test. E3 vs 1 mM Cad (p=0.0141); 0.1 mM vs 1 mM Cad (p=0.0016)

**Supplemental Figure 2.**
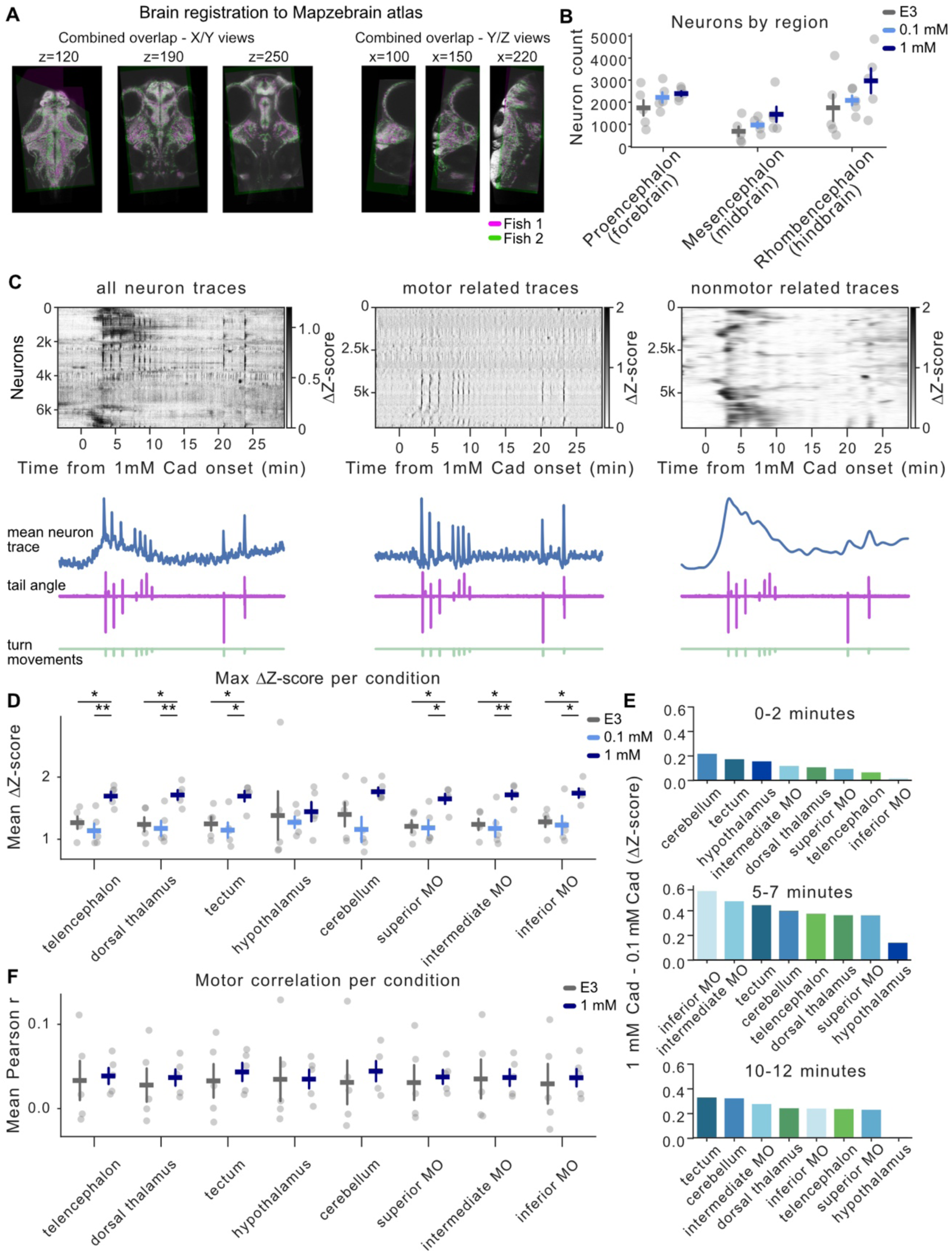
(**A**) Fish brain registration of multiple fish to the MapZeBrain atlas to compare individual neurons and regions across individuals. (**B**) Average number of neurons per brain region across fish and conditions. (**C**) Example fish given 1 mM cad showing Butterworth filtered slow and fast frequency components of neural traces and aligned tail movement. (**D**) Maximum changes in Z-score relative to baseline across regions for E3, 0.1 mM or 1 mM cadaverine conditions. Error bars denote SEM, * p<0.05, ** p<0.01. Each condition was tested for normality (Shapiro-Wilk) and analyzed using a One-way ANOVA or Kruskal-Wallis test, followed by Tukey’s or Dunn’s post hoc test with Holm correction for multiple comparison. Telencephalon (ANOVA F=9.0246, p=0.0041; 1 mM vs E3 p=0.0225; 1 mM vs 0.1 mM p=0.0042); dorsal thalamus (ANOVA F=7.6271, p=0.0073; 1 mM vs E3 p=0.0214; 1 mM vs 0.1 mM p=0.0098); tectum (Kruskal-Wallis H=8.0600, p=0.0178; 1 mM vs E3 p=0.0473; 1 mM vs 0.1 mM p=0.0267); superior MO (ANOVA F=6.6716, p=0.0113; 1 mM vs E3 p=0.0242; 1 mM vs 0.1 mM p=0.0179), intermediate MO (ANOVA F=8.3082, p=0.0054; 1 mM vs E3 p=0.0169; 1 mM vs 0.1 mM p=0.0074), inferior MO (ANOVA F=6.8937, p=0.0102; 1 mM vs E3 p=0.0263; 1 mM vs 0.1 mM p=0.0074). (**E**) Difference in Z-scored neural activity of brain regions between 1 mM cadaverine and 0.1 mM cadaverine for each time period. (**F**) Motor-related coefficients of neurons per brain region between E3 and 1mM cadaverine. No significant differences in neuron coefficients per region.

**Supplemental Figure 3.**
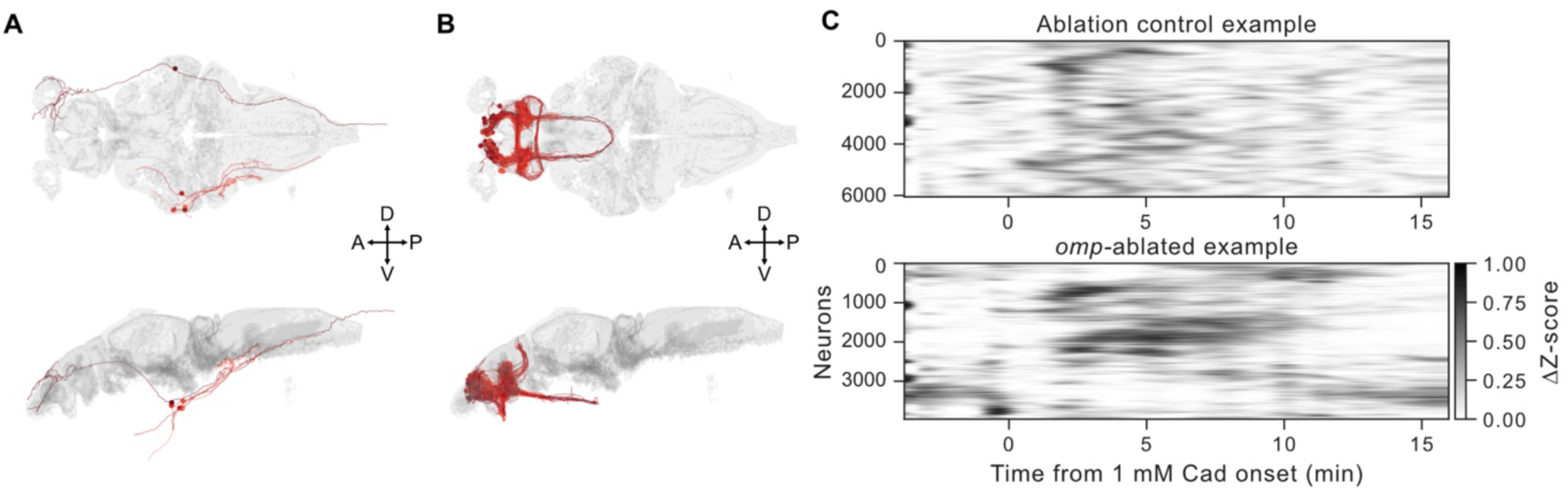
(**A**) MapZeBrain trigeminal neuron projections (**B**) MapZeBrain olfactory bulb neurons projecting to other regions of the zebrafish brain atlas. (**C**) Representative rastermaps of slow frequency neural activity of fish in ablation control and *omp* ablated fish after exposure to 1 mM cadaverine.

**Supplemental Figure 4.**
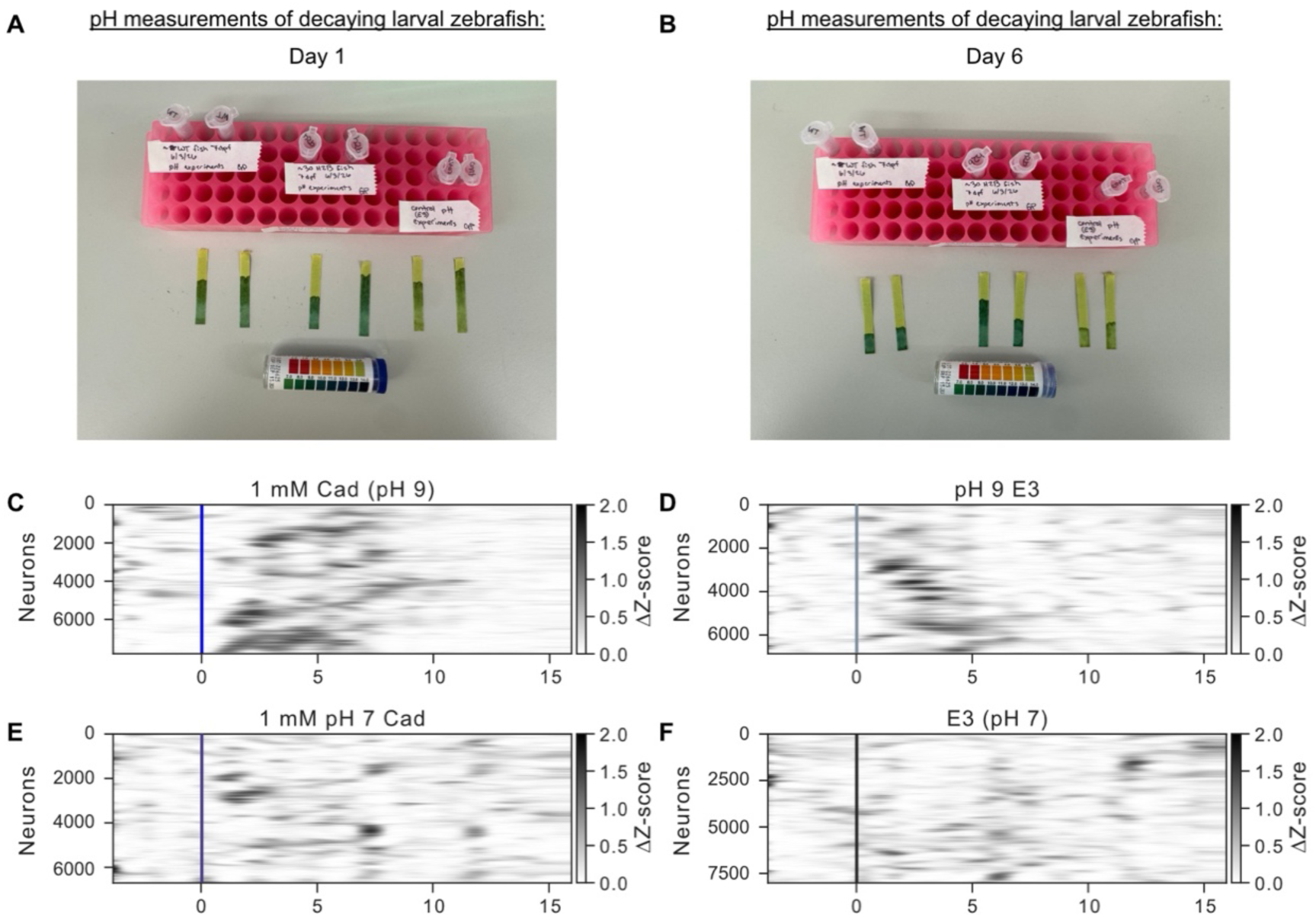
(**A**) pH of decaying larval zebrafish bodies (n=∼30 per tube) in 1 mL of E3 24 hours after death. pH was measured in tubes of either no fish, Wild type or *Tg(elavl3:H2B-GCaMP6s).* (**B**) pH of decaying zebrafish (n=∼30 per tube) in 1 mL of E3 6 days after death. pH was measured in tubes of either no fish, Wild type or *Tg(elavl3:H2B-GCaMP6s).* (**C**) Rastermap of neural activity for 1 mM cadaverine (pH 9). (**D**) Rastermap of neural activity for pH-adjusted E3 (pH 9). (**E**) Rastermap of neural activity for pH-adjusted 1 mM cadaverine (pH 7). (**F**) Rastermap of neural activity for E3 (pH 7)

**Supplemental Figure 5.**
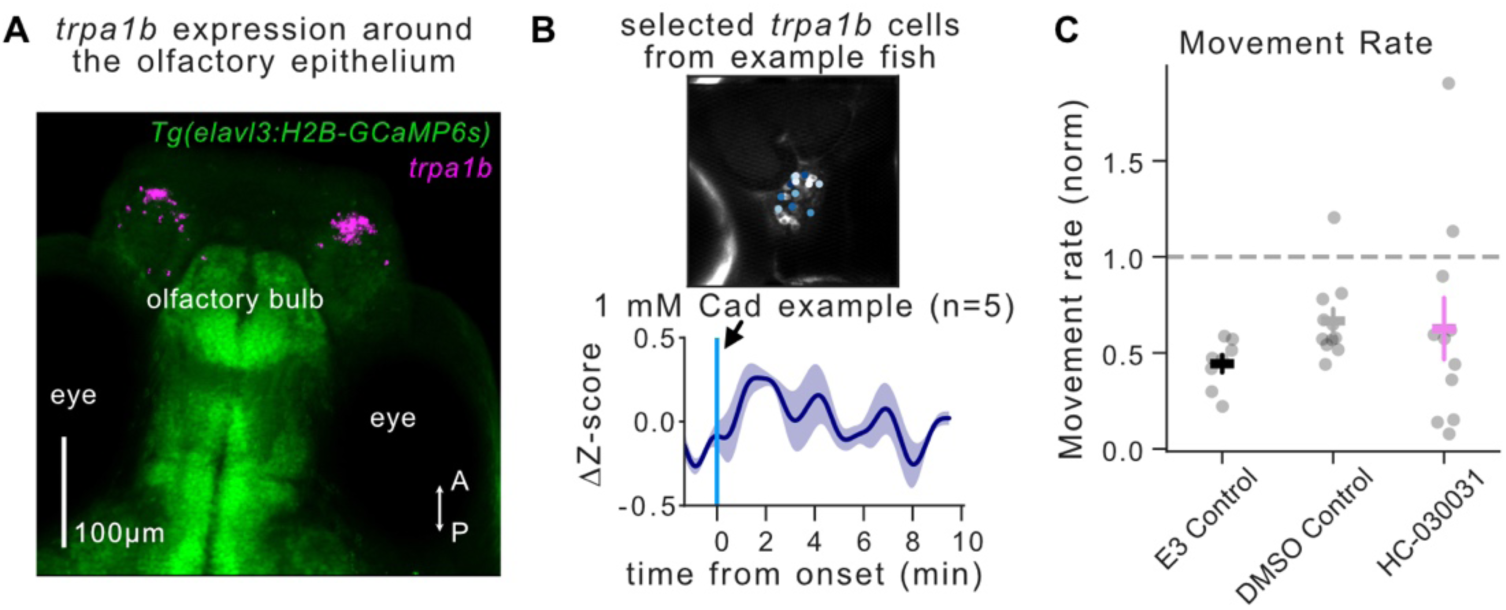
(**A**) *in situ* HCR of *trpa1b* in the forebrain, from the same fish in Fig 5A. (**B**) Max projection image of **trpa1b+** neurons in the trigeminal ganglia of *Tg(trpa1b:GCaMP6s)* fish showing ROI detection in centroid (n=12). The mean trace (shaded region denotes SEM) shows activity of cells in ROI responsive to 1 mM Cad from example fish. (**C**) Normalized movement rate of 20 trials post odor-onset compared to 5 trials pre-odor. Error bars denote SEM. One-way ANOVA revealed no significance detected between any conditions (E3 Control n=9; DMSO Control n = 10; HC-030031 n=11).

**Supplemental Figure 6.**
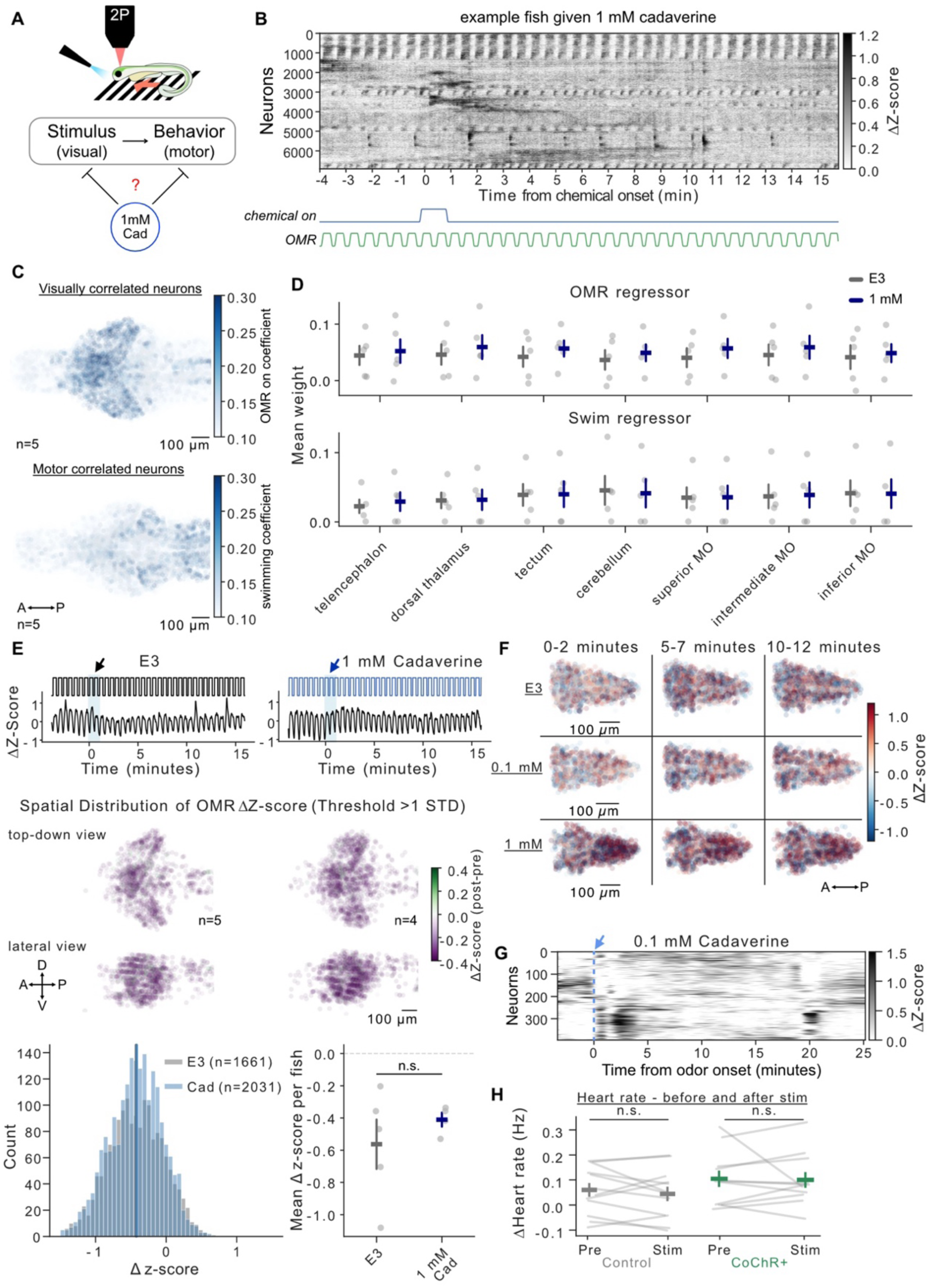
(**A**) Schematic of OMR behavior under the 2-photon microscope. Schematic illustrating the hypothesis that cadaverine modulates behavior at the level of sensory processing, motor output, or both. (**B**) Example Rastermap of calcium transients in one fish given 1 mM cadaverine while being presented closed-loop drifting gratings in a trial structure. Regressors for when the odor is on, the closed loop gratings are on/off are plotted below. (**C**) Model-derived ß-weights and location of neurons associated with visual-processing of gratings or swimming for example fish are plotted below. (**D**) OMR and swim regressors plotted across brain regions by condition (E3 or 1 mM Cad). One-way ANOVA tests found no differences between conditions or brain regions. (**E**) Example traces of visually-responsive neurons in 1 mM cadaverine or E3 trials. Spatial distribution plots of neurons and their change in z-score post-stimulus for all fish in 1 mM cadaverine (right) or E3 (left) conditions. Histogram of Δz-scores for data shown in distribution plots. Mean ΔZ-score for each condition, error bars denote SEM, and no identified significance between groups. (**F**) Spatial location and activity of GABAergic during the 0-2, 5-7, and 10-12 minutes after 1 minute exposure to E3. 0.1 mM or 1 mM Cad. (**G**) Example of rastermap showing slow-frequency responses in hindbrain gad1b neurons after exposure to 0.1 mM cadaverine. (**H**) Heart rate of *Tg(gad1b:Gal4; UAS:CoChR-eGFP)* GFP+ and GFP-fish heart rate prior to and during stimulation. The Wilcoxon tests for GFP+ fish and Paired t test on controls revealed no significance within groups.

## Notes

### Competing Interest Statement

The authors have declared no competing interest.

## References

1. Oteiza, P. & Baldwin, M. W. Evolution of sensory systems. Curr. Opin. Neurobiol. 71, 52– 59 (2021).

2. Valencia-Montoya, W. A., Pierce, N. E. & Bellono, N. W. Evolution of sensory receptors. Annu. Rev. Cell Dev. Biol. 40, 353–379 (2024).

3. v. Frisch, K. Zur Psychologie des Fisch-Schwarmes. Sci. Nat. 26, 601–606 (1938).

4. Kennedy, A. et al. Stimulus-specific hypothalamic encoding of a persistent defensive state. Nature 586, 730–734 (2020).

5. Anderson, D. J. & Adolphs, R. A framework for studying emotions across species. Cell 157, 187–200 (2014).

6. Asahina, K. et al. Tachykinin-expressing neurons control male-specific aggressive arousal in Drosophila. Cell 156, 221–235 (2014).

7. Anderson, D. J. Circuit modules linking internal states and social behaviour in flies and mice. Nat. Rev. Neurosci. 17, 692–704 (2016).

8. Lovett-Barron, M. et al. Ancestral Circuits for the Coordinated Modulation of Brain State. Cell 171, 1411–1423.e17 (2017).

9. Andalman, A. S. et al. Neuronal dynamics regulating brain and behavioral state transitions. Cell 177, 970–985.e20 (2019).

10. Mu, Y. et al. Glia accumulate evidence that actions are futile and suppress unsuccessful behavior. Cell 178, 27–43.e19 (2019).

11. Marques, J. C., Li, M., Schaak, D., Robson, D. N. & Li, J. M. Internal state dynamics shape brainwide activity and foraging behaviour. Nature 577, 239–243 (2020).

12. Flavell, S. W., Gogolla, N., Lovett-Barron, M. & Zelikowsky, M. The emergence and influence of internal states. Neuron 110, 2545–2570 (2022).

13. Atanas, A. A. et al. Brain-wide representations of behavior spanning multiple timescales and states in C. elegans. Cell 186, 4134–4151.e31 (2023).

14. Mathuru, A. S. et al. Chondroitin fragments are odorants that trigger fear behavior in fish. Curr. Biol. 22, 538–544 (2012).

15. Jesuthasan, S., Krishnan, S., Cheng, R.-K. & Mathuru, A. Neural correlates of state transitions elicited by a chemosensory danger cue. Prog. Neuropsychopharmacol. Biol. Psychiatry 110110 (2020).

16. Masuda, M. et al. Identification of olfactory alarm substances in zebrafish. Curr. Biol. 34, 1377–1389.e7 (2024).

17. Allen, C. & Hauser, M. D. Concept attribution in nonhuman animals: Theoretical and methodological problems in ascribing complex mental processes. Philos. Sci. 58, 221–240 (1991).

18. Yao, M. et al. The ancient chemistry of avoiding risks of predation and disease. Evol. Biol. 36, 267–281 (2009).

19. Choe, D.-H., Millar, J. G. & Rust, M. K. Chemical signals associated with life inhibit necrophoresis in Argentine ants. Proc. Natl. Acad. Sci. U. S. A. 106, 8251–8255 (2009).

20. Chakraborty, T. S. et al. Sensory perception of dead conspecifics induces aversive cues and modulates lifespan through serotonin in Drosophila. Nat. Commun. 10, 2365 (2019).

21. Iglesias, T. L., McElreath, R. & Patricelli, G. L. Western scrub-jay funerals: cacophonous aggregations in response to dead conspecifics. Anim. Behav. 84, 1103–1111 (2012).

22. Oliveira, T. A. et al. Death-associated odors induce stress in zebrafish. Horm. Behav. 65, 340–344 (2014).

23. Hussain, A. et al. High-affinity olfactory receptor for the death-associated odor cadaverine. Proc. Natl. Acad. Sci. U. S. A. 110, 19579–19584 (2013).

24. Kikuchi, Y., Kojima, H., Tanaka, T., Takatsuka, Y. & Kamio, Y. Characterization of a second lysine decarboxylase isolated from Escherichia coli. J. Bacteriol. 179, 4486–4492 (1997).

25. Sabo, D. L., Boeker, E. A., Byers, B., Waron, H. & Fischer, E. H. Purification and physical properties of inducible Escherichia coli lysine decarboxylase. Biochemistry 13, 662–670 (1974).

26. Dewan, A., Pacifico, R., Zhan, R., Rinberg, D. & Bozza, T. Non-redundant coding of aversive odours in the main olfactory pathway. Nature 497, 486–489 (2013).

27. Qiu, Q., Wu, Y., Ma, L. & Yu, C. R. Encoding innately recognized odors via a generalized population code. Curr. Biol. 31, 1813–1825.e4 (2021).

28. Izquierdo, C., Gómez-Tamayo, J. C., Nebel, J.-C., Pardo, L. & Gonzalez, A. Identifying human diamine sensors for death related putrescine and cadaverine molecules. PLoS Comput. Biol. 14, e1005945 (2018).

29. Dolezalova, H., Stepita-Klauco, M. & Seiler, N. Determination of cadaverine in active and dormant snails (Helix pomatia). Brain Res. 67, 349–351 (1974).

30. Brand, G. Olfactory/trigeminal interactions in nasal chemoreception. Neurosci. Biobehav. Rev. 30, 908–917 (2006).

31. Galliot, E., Laurent, L., Hacquemand, R., Pourié, G. & Millot, J.-L. Fear-like behavioral responses in mice in different odorant environments: Trigeminal versus olfactory mediation under low doses. Behav. Processes 90, 161–166 (2012).

32. Desban, L. et al. Lateral line hair cells integrate mechanical and chemical cues to orient navigation. bioRxiv 2022.08.31.505989 (2022) doi:10.1101/2022.08.31.505989.

33. Sy, S. K. H. et al. An optofluidic platform for interrogating chemosensory behavior and brainwide neural representation in larval zebrafish. Nat. Commun. 14, 227 (2023).

34. Sy, S. K. H. et al. Pallium-encoded valence-specific chemosensory amplification of eye-body coordination in larval zebrafish. eLife (2026) doi:10.7554/elife.109494.1.

35. Jenkins, B. & Frank, T. Distributed representations of chemosensory valence in a naive vertebrate brain. bioRxiv 2026.06.17.732810 (2026) doi:10.64898/2026.06.17.732810.

36. Herrera, K. J., Panier, T., Guggiana-Nilo, D. & Engert, F. Larval zebrafish use olfactory detection of sodium and chloride to avoid salt water. Curr. Biol. 31, 782–793.e3 (2021).

37. Mayseless, O. et al. Odor preference maps to cohesive transcriptional domains in the olfactory bulb. bioRxiv 2026.06.11.731560 (2026) doi:10.64898/2026.06.11.731560.

38. Wisman, A. & Shrira, I. The smell of death: evidence that putrescine elicits threat management mechanisms. Front. Psychol. 6, 1274 (2015).

39. Anderson, J. R., Yeow, H. & Hirata, S. Putrescine--a chemical cue of death-is aversive to chimpanzees. Behav. Processes 193, 104538 (2021).

40. Lovett-Barron, M. et al. Multiple convergent hypothalamus-brainstem circuits drive defensive behavior. Nat. Neurosci. (2020) doi:10.1038/s41593-020-0655-1.

41. Herrera, K. J., Zarghani-Shiraz, A., Ahrens, M. B., Engert, F. & Fishman, M. C. Synchronization of behavioral and cardiac dynamics in larval zebrafish. Cell Rep. 45, 116947 (2026).

42. Naumann, E. A. et al. From whole-brain data to functional circuit models: The zebrafish optomotor response. Cell 167, 947–960.e20 (2016).

43. Portugues, R. & Engert, F. Adaptive locomotor behavior in larval zebrafish. Front. Syst. Neurosci. 5, 72 (2011).

44. Ahrens, M. B. et al. Brain-wide neuronal dynamics during motor adaptation in zebrafish. Nature 485, 471–477 (2012).

45. Yang, E. et al. A brainstem integrator for self-location memory and positional homeostasis in zebrafish. Cell 185, 5011–5027.e20 (2022).

46. Kunst, M. et al. A cellular-resolution atlas of the larval zebrafish brain. Neuron 103, 21–38.e5 (2019).

47. Shainer, I. et al. A single-cell resolution gene expression atlas of the larval zebrafish brain. Sci. Adv. 9, eade9909 (2023).

48. Choi, J.-H., Duboue, E. R., Macurak, M., Chanchu, J.-M. & Halpern, M. E. Specialized neurons in the right habenula mediate response to aversive olfactory cues. Elife 10, (2021).

49. Dieris, M., Ahuja, G., Krishna, V. & Korsching, S. I. A single identified glomerulus in the zebrafish olfactory bulb carries the high-affinity response to death-associated odor cadaverine. Sci. Rep. 7, 40892 (2017).

50. Vendrell-Llopis, N. & Yaksi, E. Evolutionary conserved brainstem circuits encode category, concentration and mixtures of taste. Sci. Rep. 5, 17825 (2015).

51. Purves, D. et al. Trigeminal Chemoreception. in Neuroscience. 2nd edition (Sinauer Associates, 2001).

52. Haesemeyer, M., Robson, D. N., Li, J. M., Schier, A. F. & Engert, F. A Brain-wide Circuit Model of Heat-Evoked Swimming Behavior in Larval Zebrafish. Neuron 98, 817–831.e6 (2018).

53. Koide, T., Yabuki, Y. & Yoshihara, Y. Terminal nerve GnRH3 neurons mediate slow avoidance of carbon dioxide in larval zebrafish. Cell Rep. 22, 1115–1123 (2018).

54. Prober, D. A. et al. Zebrafish TRPA1 channels are required for chemosensation but not for thermosensation or mechanosensory hair cell function. J. Neurosci. 28, 10102–10110 (2008).

55. Mettam, J. J., McCrohan, C. R. & Sneddon, L. U. Characterisation of chemosensory trigeminal receptors in the rainbow trout, Oncorhynchus mykiss: responses to chemical irritants and carbon dioxide. J. Exp. Biol. 215, 685–693 (2012).

56. Sagasti, A., Guido, M. R., Raible, D. W. & Schier, A. F. Repulsive interactions shape the morphologies and functional arrangement of zebrafish peripheral sensory arbors. Curr. Biol. 15, 804–814 (2005).

57. Rosales, K., et al. Neural pathways linking hypoxia with pectoral fin movements in Danio rerio. bioRxiv (2019) doi:10.1101/655084.

58. Cain, W. S. & Murphy, C. L. Interaction between chemoreceptive modalities of odour and irritation. Nature 284, 255–257 (1980).

59. Miyasaka, N. et al. Olfactory projectome in the zebrafish forebrain revealed by genetic single-neuron labelling. Nat. Commun. 5, 3639 (2014).

60. Gau, P. et al. The zebrafish ortholog of TRPV1 is required for heat-induced locomotion. J. Neurosci. 33, 5249–5260 (2013).

61. Balakrishnan, K. A. & Haesemeyer, M. Behavioral and circuit principles of temperature gradient navigation. Curr. Biol. 35, 5395–5410.e8 (2025).

62. Lecamp, B. et al. The pro-but not antinociceptive effects of cannabidiol depend on Trpa1b in larval zebrafish. J. Neurosci. 46, e1991252026 (2026).

63. Yang, J. et al. Water volume influences antibiotic resistomes and microbiomes during fish corpse decomposition. Sci. Total Environ. 789, 147977 (2021).

64. Clements, T., Purnell, M. A. & Gabbott, S. Experimental analysis of organ decay and pH gradients within a carcass and the implications for phosphatization of soft tissues. Palaeontology 65, e12617 (2022).

65. Fujita, F. et al. Intracellular alkalization causes pain sensation through activation of TRPA1 in mice. J. Clin. Invest. 118, 4049–4057 (2008).

66. Dhaka, A. et al. TRPV1 is activated by both acidic and basic pH. J. Neurosci. 29, 153–158 (2009).

67. Roberts, A. C. et al. Induction of short-term sensitization by an aversive chemical stimulus in zebrafish larvae. eNeuro 7, ENEURO.0336-19.2020 (2020).

68. Baier, H. & Scott, E. K. The Visual Systems of Zebrafish. Annu. Rev. Neurosci. (2024) doi:10.1146/annurev-neuro-111020-104854.

69. Kramer, A., Wu, Y., Baier, H. & Kubo, F. Neuronal Architecture of a Visual Center that Processes Optic Flow. Neuron 103, 118–132.e7 (2019).

70. Chen, A. B. et al. Norepinephrine changes behavioral state through astroglial purinergic signaling. Science 388, 769–775 (2025).

71. Liberles, S. D. Trace amine-associated receptors: ligands, neural circuits, and behaviors. Curr. Opin. Neurobiol. 34, 1–7 (2015).

72. Li, Q. et al. Non-classical amine recognition evolved in a large clade of olfactory receptors. Elife 4, e10441 (2015).

73. Batalla, M., Martínez-Artero, J. & Catalan, J. Global basin-scale mapping of pH and alkalinity in inland waters. Sci. Data 13, 686 (2026).

74. Sundin, J. et al. On the Observation of Wild Zebrafish (Danio rerio) in India. Zebrafish 16, 546–553 (2019).

75. Viana, F. Chemosensory properties of the trigeminal system. ACS Chem. Neurosci. 2, 38– 50 (2011).

76. Hummel, T. & Kobal, G. Differences in human evoked potentials related to olfactory or trigeminal chemosensory activation. Electroencephalogr. Clin. Neurophysiol. 84, 84–89 (1992).

77. Zhang, J. et al. Sour sensing from the tongue to the brain. Cell 179, 392–402.e15 (2019).

78. Ye, L. et al. Enteroendocrine cells sense bacterial tryptophan catabolites to activate enteric and vagal neuronal pathways. Cell Host Microbe 29, 179–196.e9 (2021).

79. Peloggia, J. et al. Paired and solitary ionocytes in the zebrafish olfactory epithelium. Chem. Senses 50, bjaf031 (2025).

80. Takaoka, M. et al. Single-cell RNA-sequencing of zebrafish olfactory epithelium identifies odor-responsive candidate olfactory receptors. Genes Cells 30, e13191 (2025).

81. Paulsen, C. E., Armache, J.-P., Gao, Y., Cheng, Y. & Julius, D. Structure of the TRPA1 ion channel suggests regulatory mechanisms. Nature 520, 511–517 (2015).

82. Gracheva, E. O. et al. Molecular basis of infrared detection by snakes. Nature 464, 1006– 1011 (2010).

83. Valencia-Montoya, W. A. et al. Infrared radiation is an ancient pollination signal. Science 390, 1164–1170 (2025).

84. Bautista, D. M. et al. TRPA1 mediates the inflammatory actions of environmental irritants and proalgesic agents. Cell 124, 1269–1282 (2006).

85. Jordt, S.-E. et al. Mustard oils and cannabinoids excite sensory nerve fibres through the TRP channel ANKTM1. Nature 427, 260–265 (2004).

86. Bautista, D. M. et al. Pungent products from garlic activate the sensory ion channel TRPA1. Proc. Natl. Acad. Sci. U. S. A. 102, 12248–12252 (2005).

87. Bellono, N. W., Kammel, L. G., Zimmerman, A. L. & Oancea, E. UV light phototransduction activates transient receptor potential A1 ion channels in human melanocytes. Proc. Natl. Acad. Sci. U. S. A. 110, 2383–2388 (2013).

88. Bang, S. & Hwang, S. W. Polymodal ligand sensitivity of TRPA1 and its modes of interactions. J. Gen. Physiol. 133, 257–262 (2009).

89. Michel, W. C., Sanderson, M. J., Olson, J. K. & Lipschitz, D. L. Evidence of a novel transduction pathway mediating detection of polyamines by the zebrafish olfactory system. J. Exp. Biol. 206, 1697–1706 (2003).

90. Tian, L. et al. Vertebrate OTOP1 is also an alkali-activated channel. Nat. Commun. 14, 26 (2023).

91. Kang, K. et al. Analysis of Drosophila TRPA1 reveals an ancient origin for human chemical nociception. Nature 464, 597–600 (2010).

92. Major, G. & Tank, D. Persistent neural activity: prevalence and mechanisms. Curr. Opin. Neurobiol. 14, 675–684 (2004).

93. Nguyen, M. Q., Wu, Y., Bonilla, L. S., von Buchholtz, L. J. & Ryba, N. J. P. Diversity amongst trigeminal neurons revealed by high throughput single cell sequencing. PLoS One 12, e0185543 (2017).

94. Zhang, S. X. et al. Stochastic neuropeptide signals compete to calibrate the rate of satiation. Nature 637, 137–144 (2025).

95. Thornquist, S. C., Pitsch, M. J., Auth, C. S. & Crickmore, M. A. Biochemical evidence accumulates across neurons to drive a network-level eruption. Mol. Cell 81, 675–690.e8 (2021).

96. Marder, E. Neuromodulation of neuronal circuits: back to the future. Neuron 76, 1–11 (2012).

97. Vijayan, V. et al. A rise-to-threshold process for a relative-value decision. Nature 619, 563– 571 (2023).

98. Jung, Y. et al. Neurons that Function within an Integrator to Promote a Persistent Behavioral State in Drosophila. Neuron 105, 322–333.e5 (2020).

99. Vishwanathan, A. et al. Predicting modular functions and neural coding of behavior from a synaptic wiring diagram. Nat. Neurosci. 27, 2443–2454 (2024).

100. Hernandez-Nunez, L., et al. Emergence of functional heart-brain circuits in a vertebrate. bioRxiv (2025) doi:10.1101/2025.09.22.677693.

101. Arinel, M. et al. Gut distension evokes rapid neural dynamics in vagal and hindbrain populations of larval zebrafish. iScience 29, 116206 (2026).

102. Chen, W., et al. Whole-brain, all-optical interrogation of neuronal dynamics underlying gut interoception in zebrafish. bioRxiv (2025) doi:10.1101/2025.03.26.645305.

103. Freeman, J. et al. Mapping brain activity at scale with cluster computing. Nat. Methods 11, 941–950 (2014).

104. Zhang, Y. et al. Fast and sensitive GCaMP calcium indicators for imaging neural populations. Nature 615, 884–891 (2023).

105. Ferreira, T. et al. Silencing of odorant receptor genes by G protein βγ signaling ensures the expression of one odorant receptor per olfactory sensory neuron. Neuron 81, 847–859 (2014).

106. Davison, J. M. et al. Transactivation from Gal4-VP16 transgenic insertions for tissue-specific cell labeling and ablation in zebrafish. Dev. Biol. 304, 811–824 (2007).

107. Förster, D., Dal Maschio, M., Laurell, E. & Baier, H. An optogenetic toolbox for unbiased discovery of functionally connected cells in neural circuits. Nat. Commun. 8, 116 (2017).

108. Chen, T.-W. et al. Ultrasensitive fluorescent proteins for imaging neuronal activity. Nature 499, 295–300 (2013).

109. Antinucci, P. et al. A calibrated optogenetic toolbox of stable zebrafish opsin lines. Elife 9, (2020).

110. Wang, Y. et al. EASI-FISH for thick tissue defines lateral hypothalamus spatio-molecular organization. Cell 184, 6361–6377.e24 (2021).

111. Legorreta, E. M. et al. Whole-Brain Co-Mapping of Gene Expression and Neuronal Activity at Cellular Resolution in Behaving Zebrafish. bioRxiv 2026.02.07.704095 (2026) doi:10.64898/2026.02.07.704095.

112. Pachitariu, M., et al. Suite2p: beyond 10,000 neurons with standard two-photon microscopy. bioRxiv (2016) doi:10.1101/061507.

113. Lopes, G. et al. Bonsai: an event-based framework for processing and controlling data streams. Front. Neuroinform. 9, 7 (2015).

114. Guilbeault, N. C., Guerguiev, J., Martin, M., Tate, I. & Thiele, T. R. BonZeb: open-source, modular software tools for high-resolution zebrafish tracking and analysis. Sci. Rep. 11, 1– 21 (2021).

115. Lopes, G. et al. Creating and controlling visual environments using BonVision. Elife 10, e65541 (2021).

116. Zada, D. et al. Development of neural circuits for social motion perception in schooling fish. Curr. Biol. 34, 3380–3391.e5 (2024).

117. Choi, H. M. T. et al. Third-generation in situ hybridization chain reaction: multiplexed, quantitative, sensitive, versatile, robust. Development 145, dev165753 (2018).

118. Pachitariu, M. & Stringer, C. Cellpose 2.0: how to train your own model. Nat. Methods 19, 1634–1641 (2022).

119. Stringer, C. et al. Rastermap: a discovery method for neural population recordings. Nat. Neurosci. 28, 201–212 (2025).

